# Copper Acts Synergistically with Fluconazole in *Candida glabrata* by Compromising Drug Efflux, Sterol Metabolism, and Zinc Homeostasis

**DOI:** 10.1101/2021.12.22.473865

**Authors:** Ana Gaspar-Cordeiro, Catarina Amaral, Vânia Pobre, Wilson Antunes, Ana Petronilho, Paulo Paixão, António P. Matos, Catarina Pimentel

## Abstract

The synergistic combinations of drugs are promising strategies to boost the effectiveness of current antifungals and thus prevent the emergence of resistance.

In this work, we show that copper and the antifungal fluconazole act synergistically against *Candida glabrata*, an opportunistic pathogenic yeast intrinsically tolerant to fluconazole.

Analyses of the transcriptomic profile of *C. glabrata* after the combination of copper and fluconazole showed that the expression of the multidrug transporter gene *CDR1* was decreased, suggesting that fluconazole efflux could be affected. In agreement, we observed that copper inhibits the transactivation of Pdr1, the transcription regulator of multidrug transporters and leads to the intracellular accumulation of fluconazole. Copper also decreases the transcriptional induction of ergosterol biosynthesis (ERG) genes by fluconazole, which culminates in the accumulation of toxic sterols. Co-treatment of cells with copper and fluconazole should affect the function of proteins located in the plasma membrane, as several ultrastructural alterations, including irregular cell wall and plasma membrane and loss of cell wall integrity, were observed. Finally, we show that the combination of copper and fluconazole downregulates the expression of the gene encoding the zinc-responsive transcription regulator Zap1, which possibly, together with the membrane transporters malfunction, generates zinc depletion. Supplementation with zinc reverts the toxic effect of combining copper with fluconazole, underscoring the importance of this metal in the observed synergistic effect.

Overall, this work, while unveiling the molecular basis that supports the use of copper to enhance the effectiveness of fluconazole, paves the way for the development of new metal-based antifungal strategies.

## 1. Introduction

Invasive fungal infections (IFIs) are a major health concern, affecting many people worldwide and causing more than 1.2 million deaths per year (1). IFIs caused by the yeast Candida, also named invasive candidiasis, represent the most common fungal disease among hospitalized patients receiving immunosuppressive or intensive antibacterial therapies (2, 3). Candida blood stream infections, known as candidemias, are the prevalent form of invasive *c*andidiasis and a serious problem in intensive care units, where between half to two-thirds of these episodes occur (3, 4). *Candida albicans* is the predominant causative organism of candidemia, but recent data have shown a marked epidemiologic shift toward non-albicans species, such as *Candida glabrata* and *Candida parapsilosis*, which are more resistant to antifungals (5).

Current invasive candidiasis treatment guidelines include the antifungal fluconazole as a primary therapeutic option (6). Fluconazole acts by inhibiting lanosterol 14-α-demethylase (Erg11), a heme-containing cytochrome P450 enzyme, involved in the biosynthesis of ergosterol, an important component of fungal membranes (7). Inhibition of Erg11 leads to the accumulation of toxic methylated sterols in the membrane, which arrests cell growth and division (7, 8). The fungistatic rather than the fungicidal properties of fluconazole together with its intensive use and misuse have paved the way for the appearance of resistant clinical isolates, with *Candida glabrata* standing out among them (5). In fact, invasive *Candida glabrata* infections are associated with high rates of morbidity and mortality plausibly because of the emergence of resistance to available antifungals (9, 10). Also the ability of the yeast to escape the immune system and thrive in the hostile host environment is acknowledged as contributing to that scenario (reviewed in (11)). In *C. glabrata*, the development of azole resistance has been almost exclusively associated with the presence of activating mutations in the Zn_2_Cys_6_ transcription factor Pdr1 (12, 13). These mutations lead to the overexpression of genes encoding drug efflux proteins (Cdr1 and Cdr2), which results in the decrease of fluconazole within cells (12, 14).

The antimicrobial properties of copper have been known for centuries and stem from its toxicity, when present above physiological needs (15). Copper also plays a central role at the host-pathogen axis. While from the host side, it is required for the differentiation and maturation of immune cells and is used as a weapon by phagocytes, as part of the nutritional immune response (16), from the pathogen perspective, a fine-tuned copper homeostatic machinery is critical for survival within the host (17).

Inspired by the antifungal properties of copper, several authors have combined this metal with fluconazole and showed that this could be a promising strategy, since the combination resulted in greater antifungal activity (18-20). However, the mechanism underlying such synergism has yet to be elucidated.

In this study, using a genetic and biochemical approach, we investigated how copper and fluconazole act synergistically against *Candida glabrata*. Our data indicate that an excess of copper inhibits the activity of Pdr1, which leads to the accumulation of fluconazole and causes abnormal sterol biosynthesis. In addition, we identified zinc homeostasis as a central player in copper fluconazole synergism, which puts zinc under the spotlight as a putative target for future antifungal investment.

### 2. Materials and Methods

### Strains, growth conditions

The strains used in this study are listed in Table S1. Yeast species were maintained in YPD agar plates. Unless otherwise stated, all assays were performed in SC medium pH 5.5 (Complete Supplement Mixture: 0.77 g/L; yeast nitrogen base w/o amino acids and ammonium sulphate 1.7 g/L; ammonium sulphate 5.4 g/L; glucose 2%). Unless otherwise stated, yeast cultures were grown until exponential phase (OD_600_ 0.8) and treated with 625 µM CuSO_4_, 32 µg/mL fluconazole or with both compounds for 24 h. For zinc supplementation assays 4 mM of ZnSO_4_ was used. Spot assays were carried out by spotting 5 µL of sequential dilutions (*Candida glabrata:* from 8×10^6^ to 80 cells/mL and *Saccharomyces cerevisiae:* 4×10^6^ to 40 cells/mL) of exponentially growing cultures onto SC agar plates containing the indicated treatments. Unless otherwise stated, the plates were incubated for 48 h at 37 ºC. Spot assays were repeated at least three times. To construct the *C. glabrata* Δ*zap1* mutant (Cg Δ*zap1*), a histidine cassette was constructed by amplifying Cg*HIS3* gene with primers for the flanking region of the Cg*ZAP1* gene (pZAP1_KO_FW: 5’CATAATTGAGTATAAAAAGGACTACAAAAGCTAAGAAACTACAGACCGGCTTGGGTACAAGAGGATAAC3’/ZAP1_KO_RV:5’GGACGAGTTGGAGTTATGTCTATATATTTATTGGACTATGTACAGTATAGTGAATGGGCAATGCTATGGTGTTGTTGAAACCAC CACGAG3’). *C. glabrata* competent cells and transformation were performed with the Zymo transformation kit (Zymo Research), according to manufacturer’s instructions. Mutant cells were confirmed by colony PCR using specific *ZAP1* primers (ZAP1_A1: 5’TGATTGTGGTGGGAGATGC3’ and ZAP1_A4: 5’GTTGATCACAGCGAGAAGC3’), followed by fragment digestion with *XcmI*. To construct the *pZAP1* plasmid, the C*gZAP1* gene was amplified by PCR with primers for the flanking region of this gene (ZAP1_A1N: 5’AATGCAAGAGGGACAAATCG3’ and ZAP1_A4). The PCR product was inserted into the *SmaI-*digested pCgACT14 vector using T4 DNA ligase (NEW ENGLAND BioLabs Inc.), according to manufacturer’s instructions.

### Pdr1 transactivation assay

The transactivation activity of *C. glabrata* Pdr1 was investigated using a *LexA*-based one-hybrid system. The *CgPDR1* ORF (CAGL0A00451g) was amplified by PCR using specific oligonucleotides (*PDR1_ATG_*II*_*FW: 5’ ATGCAAACATTAGAAACTACATCAAAATC 3’/ PDR1_KpnI_Rv: 5’ ccGGTACCTCACAAGTAAACATCAGAAAATAGGTC 3’). The PCR product was digested with *KpnI* and inserted into the *Sma*/-*KpnI* digested Yap8-LexA plasmid (21), in frame with the *LexA* sequence. The plasmid (CgPdr1-*LexA*) sequence was confirmed by sequencing. The *S. cerevisiae* EGY48 strain was then co-transformed with CgPdr1-*LexA* and pSH18-34 (ThermoFisher), which contains eight *LexA* operators fused to the *LacZ* reporter gene. Positive transformants were selected in SD agar plates, after 48 h incubation at 30ºC. For the transactivation assay, the co-transformed strain was grown in SD medium until exponential phase and left untreated or treated with 64 µg/mL of fluconazole, 625 µM of CuSO_4_ or with both compounds, for 2 h. Total proteins were extracted from cell cultures as described in (22), resolved in a 10% SDS-PAGE and immunoblotted using a monoclonal anti-β-galactosidase antibody and a monoclonal anti-Pgk1 antibody. Pgk1 was used as loading control. Immunoblots were repeated twice, with different protein extracts.

### Quantitative RT-PCR

*Candida glabrata* cultures were grown under the indicated conditions, harvested by centrifugation and RNA was isolated using the phenol:chloroform method (23). RNA samples were treated with DNase and purified using the RNeasy Kit (Qiagen). Total RNA (1 µg) was reverse transcribed with “Transcriptor reverse transcriptase” (Roche), according to the manufacturer’s instructions. qPCR reactions were performed in the Light Cycler 480 II Real-Time PCR System (Roche). Relative standard curves were constructed for each gene, using serial dilutions of cDNA. The relative expression of the genes was calculated by the relative quantification method with efficiency correction, using the LightCycler 480 Software 1.5 (Roche). The *RPL10* gene (CAGL0K12826g), which encodes an ortholog of the *S. cerevisiae* ribosomal L10 protein, was used as reference gene. All assays were performed using biological triplicates and technical duplicates. The following primer pairs were used: *CDR1*: 5’ CATGGCCACTTTTGGTCTTT 3’/5’ CAGCAATGGAGACACGCTTA 3’; *PDR1*: 5’ CAGCAATGGAGACACGCTTA 3’/5’ AGTGGGCACGTCAGAGACAG 3’; *ERG11*: 5’ TATGGTCGCCTTGCCATT 3’/5’ GACCCATGGGATCCAGTAGA 3’; *ZAP1:* 5’ GAAGCCTTACGAGTGCCATA 3’/5’ TGCACTTCAGTGGTTTCTCA 3’; *RPL10:* 5’ GAGATTCTTTCCACTTGAGAGTCAGA 3’/5’ CTCTCATACCTTGTTGCAATCTATCC 3’.

### ROS quantification

Flow cytometry analysis for ROS detection was performed as described by da Silva *et. al* (24). Cultures grown for 24 h in the presence of the indicated compounds were harvested by centrifugation, washed with PBS and resuspended in PBS containing 5 µg/mL of dihydrorhodamine 123 (DHR123). After incubation in the dark at room temperature for 2 h, cells were washed with PBS and filtered. Assays were performed in an S3e cell sorter (Bio-rad) and DHR123 signal was detected using the 488 nm excitation laser and the FL1 detection channel. Unstained cells were used to establish the gate of cells exhibiting positive DHR123 signal. A total of 100,000 gated cells were counted for each sample. Five biological replicates of each condition were analyzed and the average and SD of the mean of the FL1 (DHR123 signal) peak area was calculated.

### Cell viability assays

Cultures grown for 24 h in the presence of the indicated compounds were harvested by centrifugation, washed with PBS and resuspended in PBS containing 50 µg/mL of propidium iodide (PI). After a 15 min incubation period, samples were washed with PBS and filtered. Assays were performed in an S3e cell sorter (Bio-rad) and PI signal was detected using the 488 nm excitation laser and the FL3 detection channel. Unstained cells were used to establish the baseline of cells exhibiting positive PI signal. Five biological replicates of each condition were analyzed and a total of 100,000 cells were counted in each sample.

### Quantification of intracellular metals

Cultures were grown under the indicated conditions for 24 h, harvested and washed with 10 mM EDTA and metal-free water as described in (25). Total copper and zinc cellular contents were measured by inductively coupled plasma atomic emission spectroscopy (ICP-AES) at REQUIMTE – LAQV, Universidade Nova de Lisboa, Caparica, Portugal. Data was normalized against OD_600_. All assays were made using biological quadruplicates.

### Sterol identification and quantification by gas chromatography-mass spectrometry (GC-MS)

Sterol extraction and GC-MS analyses were performed according to the protocols described by Morio et al. and Demuyser et al. (26, 27), with minor adaptations. Briefly, cultures grown for 24 h in the presence of the indicated compounds were harvested by centrifugation. Pellets were resuspended in saponification medium (25 g of KOH, 36 mL of distilled water and made up to 100 mL with 100% ethanol), vortexed and incubated for 1 h at 80 ºC. After addition of 1 mL miliQ water, 4 mL of hexane and 50 µg of cholestane (internal standard), samples were vortexed and the hexane layer recovered. Hexane was evaporated and samples were derivatised with sylliating mixture (85432; Sigma), for 30 min. An additional 500 µL of hexane was added and samples were centrifuged at 2500 g for 5 min and supernatant was recovered. For the GC-MS analysis, 1 µL of each sample was injected into a DB5 column (5% Phenyl, 95 % Dimethyl Polysiloxane) in spitless mode using helium as carrier gas at a flow rate of 1 mL/min. The inlet was held at 200°C. The column oven was held at 50°C for 1 min, and then ramped up, at a rate of 50°C per min, to 260°C, followed by a ramp rate of 4°C per min up to the final temperature of 320°C, and then held for 5 min. The total run time was 25.2 min. Analyses were performed at REQUIMTE – LAQV, Universidade Nova de Lisbon, Caparica, Portugal. Sterols were identified according to their retention time (relative to cholestane) and by comparison with GC-MS profiles described in (28).

### Ergosterol quantification by HPLC

*C*ultures were left untreated or treated as indicated, for 24 h. Ergosterol extraction and quantification was done as described in (29), adapted from (30). Analyses were performed using a Waters 2695 Alliance HPLC system (Waters Chromatography, Milford, MA) equipped with a Waters 486 Absorbance Detector set at 282 nm. Samples (10 µL) were injected into a Symmetry C18 column (4.6 × 250 mm, 5 µm particle size, Waters Chromatography, Milford, MA), using 95% methanol as eluent in isocratic mode at a 1.1 mL/min flow rate. Column temperature was held at 30ºC and samples were kept at 10ºC. The Empower 2 software (Waters Chromatography, Milford, MA) was used for data acquisition. The retention time of the compound (15.7 min) was compared with standards for identification and the peak area was used for quantification. A standard curve prepared with commercial ergosterol was plotted to determine the amount of ergosterol present in each sample. All assays were performed using biological quadruplicates.

### Fluconazole quantification by HPLC

*Candida glabrata* cultures were grown under the indicated conditions for 24 h, harvested, resuspended in 2 mL MilliQ water and normalised against OD_600_. Tinidazole (100 µg) was added to samples prior to extraction. Extraction was then carried out using glass beads and methanol. After 30 s of vortex agitation, samples were incubated at 30ºC and 200 rpm for 1 h, followed by centrifugation at 4000 g for 7 min at 4ºC. The supernatant was then recovered and centrifuged again at 9500 g for 10 min, to remove any remaining debris. Analyses were performed using a Waters 2695 Alliance HPLC system (Waters Chromatography, Milford, MA) equipped with a Waters 486 Absorbance Detector set at 245 nm. Samples (80 µL) were injected into a reverse phase Symmetry C18 column (4.6 × 250 mm, 5µm particle size, Waters Chromatography, Milford, MA) using an isocratic eluent mixture of ammonium acetate buffer/acetonitrile/methanol (80:15:5 % v/v) at 0.7 mL/min flow rate. Column temperature was held at 30 ºC and samples were kept at 10 ºC. The Empower 2 software (Waters Chromatography, Milford, MA) was used for data acquisition. The retention time of fluconazole (13.4 min) was compared with standards for identification, and the peak area was used for quantification. Quantification of fluconazole was normalized to the internal control (tinidazole, retention time 10.4 min). Standard curves for fluconazole and tinidazole were plotted to measure the amount of compound present in each sample. All assays were performed using biological quadruplicates.

### Immunoblotting

The BVGC3 strain was grown until exponential phase (OD_600_ 0.8), treated with with 625 µM CuSO_4_, 32 µg/mL fluconazole or with both compounds for up to 2.5 h and harvested. Total proteins were extracted from cell cultures as described in (22). Protein extracts (50 µg) were resolved in a 12% SDS/PAGE gel and immunoblotted. An anti-HA-Peroxidase high affinity rat monoclonal antibody (Roche) was used to detect the HA-tagged version of Erg11 (31). Pgk1 was used as loading control and detected using a monoclonal antibody (Life Technologies). Immunoblots were repeated at least twice with different protein extracts.

### RNA-Seq and data analysis

*Candida glabrata* ATCC2001 was grown to exponential phase (OD_600_ 0.8) in SC medium and left untreated or treated with 1.25 mM CuSO_4_, 32 µg/mL fluconazole or a combination of both for 1 h. Biological triplicates were used for each condition. Cells were harvested and RNA was isolated. RNA samples were treated with DNase and purified using the RNeasy Kit (Quiagen). Library preparation and sequencing was done at the Genomics facility of Instituto Gulbenkian da Ciência, Portugal, using the SMART-SEQ2 protocol, adapted from (32), and an Illumina NextSeq500 platform (single end, 75-bp read length, 20 M reads), respectively. The quality of the RNA-Seq raw data was analysed using the FastQC tool. We mapped the reads against *Candida glabrata* genome (GCA_000002545.2 downloaded from NCBI genome database) using Bowtie2 program and obtained more than 90% of aligned reads (33). The mapping files were sorted by genomic position using Samtools and the quantification of the transcripts expression was done using the FeatureCounts software (34, 35). Differential expression analyses were done with the R package edgeR (36). We considered all transcripts with a False Discovery Rate (FDR) correction of the p-value lower than 0.05 as significant and we further filtered our results by establishing cut-offs of expression (Log_2_CPM – counts per million -higher than 3) and fold-change (FC) higher than 2. The functional annotation was performed with FungiFun (37).

### Checkerboard assays

The assays were performed in SC medium using 96-well flat bottom plates, according to the CLSI standard M27-A3 (38), following the protocol described in (39, 40). Fluconazole and copper stock solutions were prepared in bi-distilled water, filtered and further diluted in medium. Two-fold dilutions were prepared for each drug so that final concentrations ranged from 128 to 0.125 µg/mL for fluconazole and 2500 to 39 µM for CuSO_4_. A 2 × 10^6^ cells/mL suspension obtained from a single colony was diluted to 3 × 10^3^ cells/mL, in SC medium, to prepare the final inoculum. MIC endpoints were defined as the lowest drug concentration leading to a growth reduction higher than 50%, as determined by measuring OD_600_. The type of interaction for each copper and fluconazole combination tested was determined by calculating the fractional inhibitory concentration (FIC) index (ΣFIC = FIC_Cu_ + FIC_Fluc_). The MIC for fluconazole and copper were determined to be 64 µg/mL and 625 µM, respectively. For each drug combinations the FIC was calculated as follows: FIC_Cu_ = MIC_Cu+Fluc_/625 and FIC_Fluc_ = MIC_Fluc+Cu_/64. FIC index ≤ 0.5 indicates a synergistic effect.

### Scanning electron microscopy

*Candida glabrata* cultures were grown under the indicated conditions for 24 h and harvested by centrifugation. Samples were fixed with a mixture of 0.4% glutaraldehyde and 4% formaldehyde in sodium cacodylate buffer 0.1 M for 18 h at room temperature, followed by three washing steps with sodium cacodylate buffer 0.1 M. Samples were dehydrated by sequential washing with four ethanol solutions (50%, 70%, 90% and 100%). After the last dehydration step, ethanol was removed and tert-butyl alcohol was added. Samples were incubated at 30 ºC for 1 h, placed on ice until completely frozen, freeze-dried by lyophilisation and kept at room temperature until the SEM analyses were performed. Samples were gold sputtered with 8 nm gold in an electron sputter (Cressington 108) and imaged in a Hitachi SU8010 scanning electron microscope, at 1.5 kV. Approximately 120 cells were counted per condition and the proportion of cells exhibiting surface abnormalities was calculated

### Transmission electron microscopy

*Candida glabrata* cultures were grown under the indicated conditions for 24 h, harvested by centrifugation and fixed in 3% glutaraldehyde in 0.1M sodium cacodylate buffer pH 7.3. Following primary fixation for 2 h at 4ºC and wash in the cacodylate buffer, cells were, pelleted and embedded in 2% agar for further processing. Samples were further fixed for 3 h in 1% osmium tetroxide in 0.1M sodium cacodylate buffer pH 7.3. Then, samples were washed in 0.1M acetate buffer, pH 5.0 and fixed in 1% uranyl acetate in the same buffer for 1 h. Dehydration was carried out with increasing concentrations of ethanol. After passing through propylene oxide, samples were embedded in Epon-Araldite, using SPI-Pon as an Epon 812 substitute. Thin sections were made with diamond knives and stained with 2% aqueous uranyl acetate and Reynold’s lead citrate. The stained sections were analysed and photographed in a JEOL 1200-EX electron microscope

### Data availability

RNA-Sequencing raw data is available at Gene Expression Omnibus (GEO) under the accession number GSE162741.

## 3. Results

### 3.1 Copper acts synergistically with fluconazole

Growing evidence indicates that copper potentiates the antifungal activity of fluconazole against opportunistic pathogenic yeasts (18-20). Here, we confirmed that the combination of copper and fluconazole impairs the growth of *Candida glabrata* at concentrations where these compounds cause only mild growth defects when used separately (Fig. 1A). Accordingly, by flow cytometry using propidium iodide (PI), a membrane impermeable dye generally excluded from viable cells, we found that, when used together, copper and fluconazole strongly increased PI staining by more than two-fold compared with individual treatments (Fig. 1B). Noteworthy, clinical isolates and other laboratory strains exhibited a similar growth phenotype (Fig. S1 A and B), indicating that the effect is not restricted to the most used strain in this study (ATCC2001). Consistent with previous reports (41), we observed that *Candida glabrata* colonies exhibited a brown pigmentation in the presence of copper, presumably as a result of CuS mineralization on the cell surface (42). Notably, co-treatment with fluconazole decreased this pigmentation (Fig. 1A).

**FIG 1.**
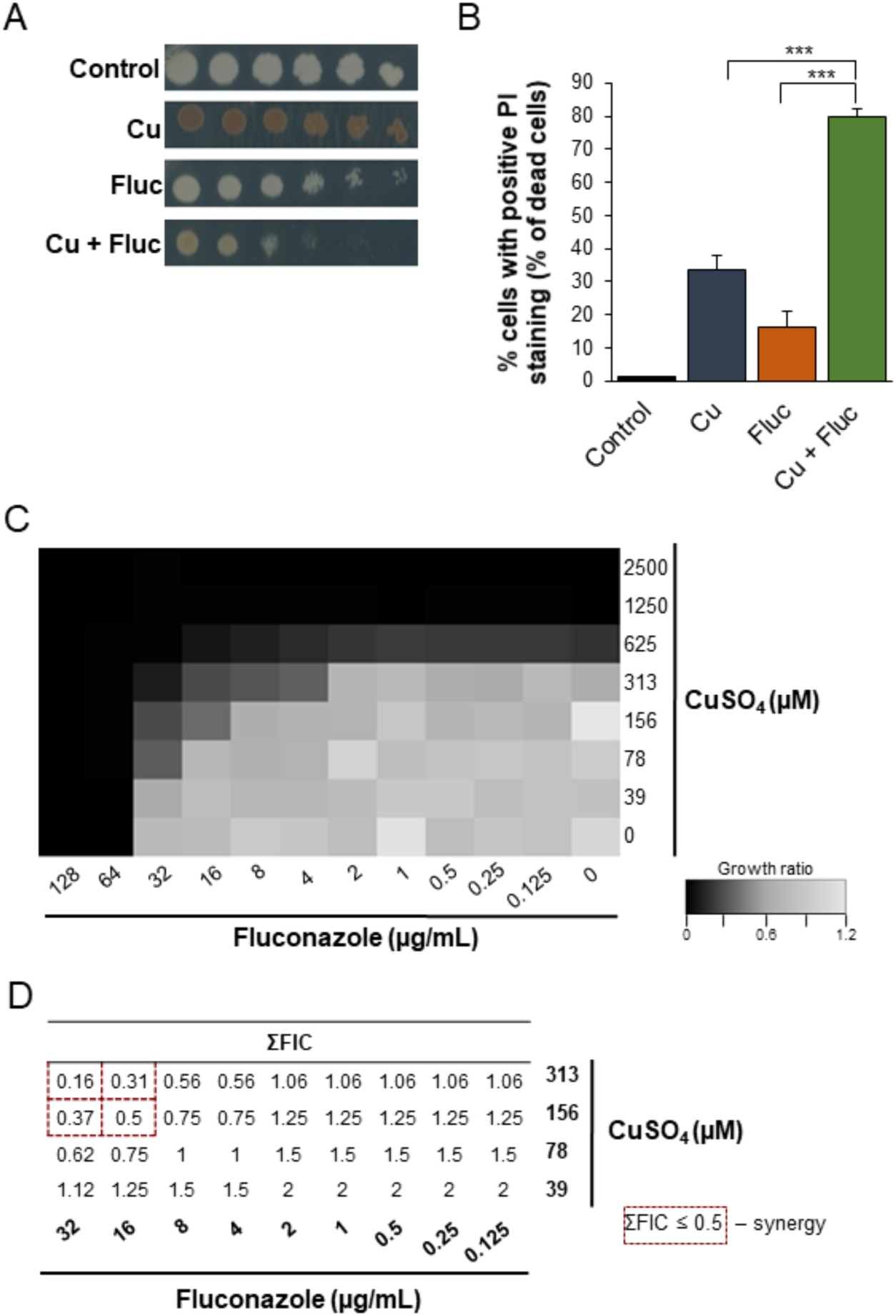
Copper acts synergistically with fluconazole. A: Growth sensitivity of *Candida glabrata* in SC agar plates (Control) containing 32 µg/mL of fluconazole (Fluc), 625 µM CuSO_4_ (Cu) or a combination of both (Cu+Fluc). Growth was recorded after 48 h at 37 ºC. B: Detection of dead cells by flow cytometry using propidium iodide staining (PI) under control conditions or after treatment with 625 µM CuSO_4_ (Cu), 32 µg/mL fluconazole (Fluc) or both (Cu+Fluc) for 24 h. Significance of differences was calculated using student’s T-test (*** p-value < 0.001). C: Checkerboard assays. Cells were exposed to all the possible combinations of CuSO_4_ and fluconazole (ranging between 2500 – 39 µM for CuSO_4_ and 128 – 0.125 µg/mL for fluconazole). Growth ratios were evaluated after 24 h at 37 ºC by measuring OD_600_ and normalizing it relative to untreated controls (see gradient bar). D: Fractional Inhibitory Concentration (FIC) index (ΣFIC) was calculated for the indicated combinations of fluconazole and CuSO_4_. The dashed red squares indicated the Cu+Fluc concentrations where synergy is observed.

The interaction between copper and fluconazole was further investigated using checkerboard microdilution assays. We tested the effect on cell growth of all possible combinations of copper and fluconazole, ranging from 2500 to 39 µM and 128 to 0.125 µg/mL, respectively. The results are summarized in Fig. 1C and D. The MIC for fluconazole (MIC_Fluc_) was 64 µg/mL and for copper (MIC_Cu_) 625 µM. Copper reduced the MIC_Fluc_ four-fold (16 µg/mL) and sixteen-fold (4 µg/mL) when added at concentrations of 156 and 313 µM, respectively. Fluconazole decreased the MIC_Cu_ four-fold (156 µM) and eight-fold (78 µM) when combined with copper at concentrations of 32 and 16 µg/mL, respectively (Fig. 1C). Synergistic effects (ΣFIC ≤ 0.5) between copper and fluconazole were observed for concentrations equal to or higher than 156 µM (CuSO_4_) and 16 µg/mL (fluconazole), as depicted in Fig. 1D. The effect was also observed in a *Candida glabrata* strain with a lower MIC for fluconazole (Cg*Δcdr1*, Fig. S1 C and D).

### 3.2 Differences in copper accumulation do not explain the synergistic effect between copper and fluconazole

Fluconazole is capable of forming complexes with metals, which likely increases its antifungal activity (18). Therefore, we first hypothesized that the observed synergism between the drug and copper (Fig. 1) could be a result of the formation of a fluconazole-copper complex. However, we were unable to observe such a complex under our experimental conditions (data not shown). Similar results were reported by Hunsaker and Franz (19), clearly showing that a more intricate relationship should underpin the observed synergistic effect. Interestingly, those authors also demonstrated that adaptation to fluconazole in *Candida albicans* required the modulation of metal homeostasis and that cells exposed to the drug accumulated more copper (43). In this sense, we started by testing if the synergy between copper and fluconazole in *Candida glabrat*a could also be due to the toxic accumulation of copper. To do so, we quantified the levels of copper in cells treated for 24 h with 32 µg/mL fluconazole, 625 µM CuSO_4_, or a combination of both compounds (hereafter referred to as Cu+Fluc). Unlike *C. albicans* (43), Cu+Fluc-treated *Candida glabrata* cells were found to accumulate far less copper when compared with those treated with the metal alone (Fig. 2A). In agreement with this observation, cells exposed to Cu+Fluc also had lower levels of ROS, a well-known mechanism of copper toxicity (15), as measured by flow cytometry using DHR123 (Fig. 2B).

**FIG 2.**
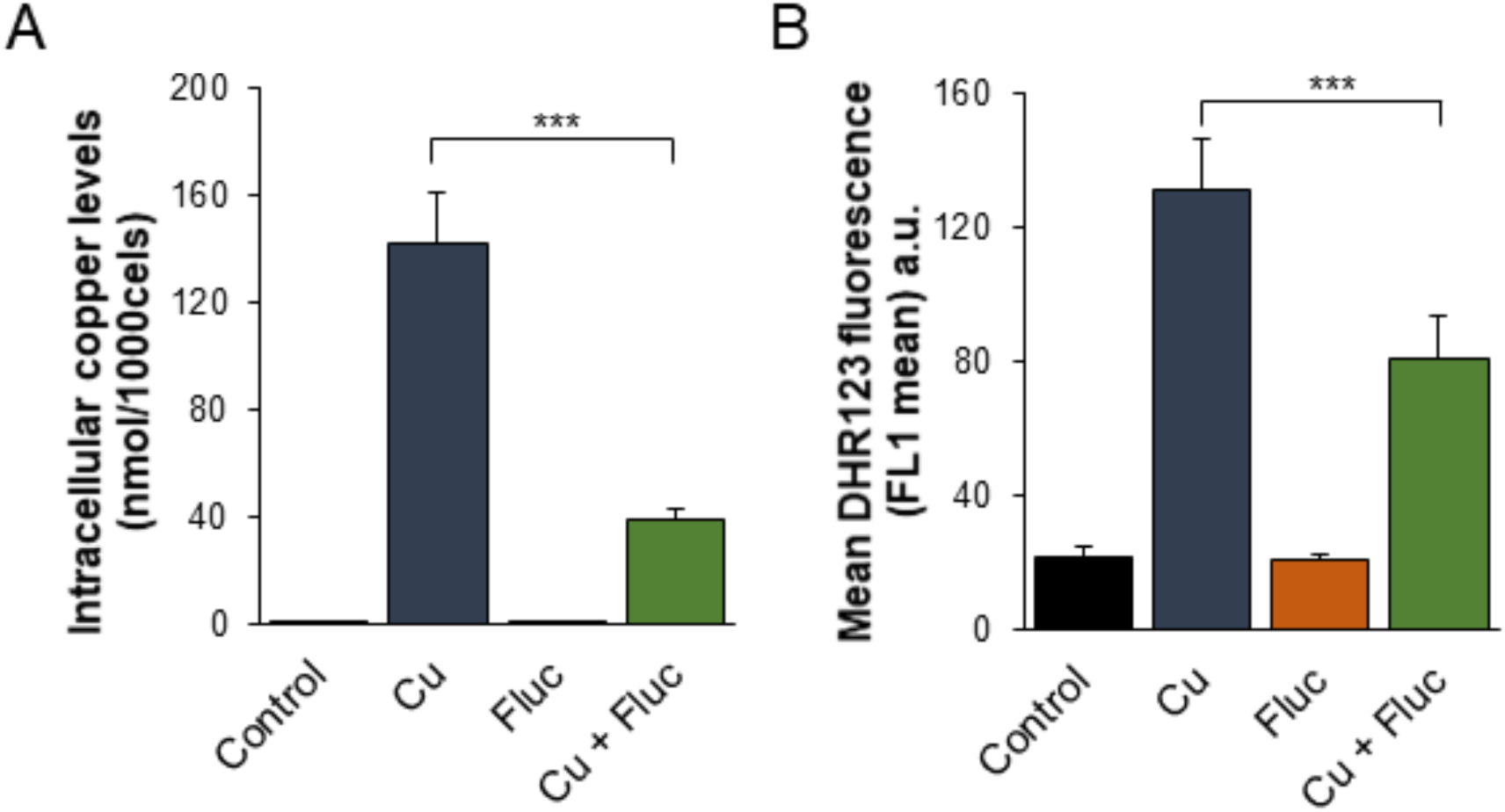
Differences in copper accumulation do not explain the synergistic effect between copper and fluconazole. A: The copper content of *Candida glabrata* cells left untreated (Control) or treated with 625 µM CuSO_4_ (Cu), 32 µg/mL fluconazole (Fluc) or both (Cu+Fluc) for 24 h was determined by ICP-AES. B: ROS detection in *Candida glabrata* cells left untreated (Control) or treated 625 µM CuSO_4_ (Cu), 32 µg/mL fluconazole (Fluc) or both (Cu+Fluc) for 24 h, was performed by flow cytometry using DHR123. Values are the mean of five independent biological replicates with 100,000 cells counted per each condition. Significance of differences was calculated using the student’s T-test (*** p-value < 0.005).

These data indicate that in *Candida glabrata*, the antifungal synergy between copper and fluconazole cannot be attributed to exacerbated copper accumulation and that other mechanism(s), potentially specific to this organism, should underlie such effect.

### 3.3 Transcriptional response of *Candida glabrata* to copper and fluconazole

In yeast species, the response to copper overload requires the transcriptional activation of copper detoxification genes, such as those encoding metallothioneins (44), by a copper-responsive transcription factor (Amt1 in *C. glabrata* (45)) The adaptation to fluconazole also relies on the transcriptional activator, Pdr1 (12, 13, 46), which is responsible for activating multidrug efflux genes, such as *CDR1* and C*DR2* (14, 46). Therefore, as a first approach to understanding the antifungal synergism between copper and fluconazole, we decided to investigate the alterations in the transcriptional landscape of *C. glabrata* exposed to Cu+Fluc and compared them with those of cells treated with fluconazole and copper separately. Differential gene expression analyses (relative to untreated cells) were conducted using edgeR (36), and a set of restrictive parameters was applied (log_2_FC > 1, log_2_CPM > 3, and FDR < 0.05) to identify the most relevant affected pathways.

After exposure to fluconazole, only 34 genes were found differentially expressed, whereas copper and Cu+Fluc altered the expression of 1164 and 1451 genes, respectively. The overlapping and specific alterations of each transcriptomic profile are illustrated in Fig. 3A and the genes whose expression was affected by the individual or combined treatments are listed in Tables S2–S4.

**FIG 3.**
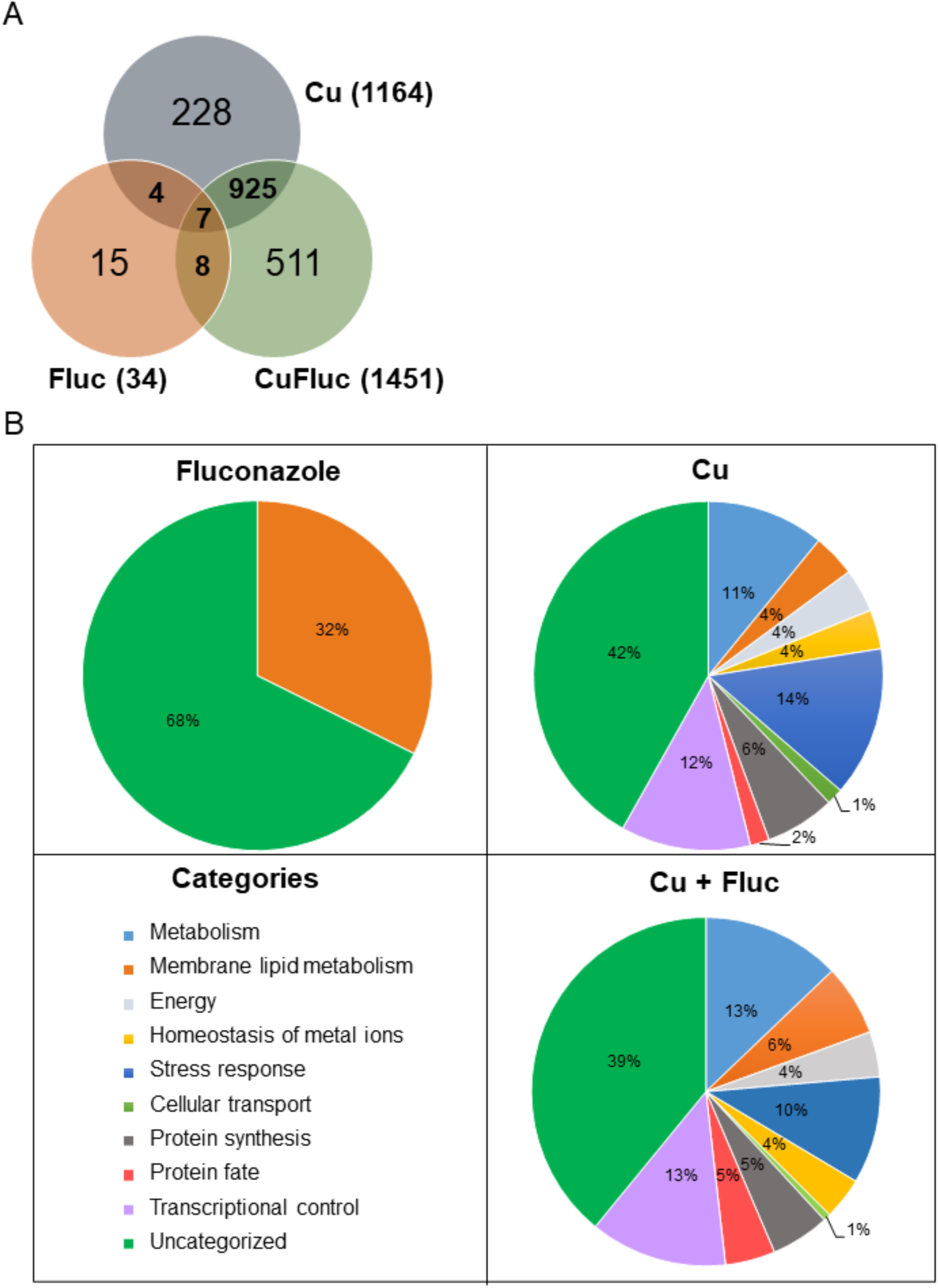
Transcriptional response *of Candida glabrata* to copper and fluconazole. A: Venn diagram representing the number of genes differentially expressed. The total number of differentially expressed genes in each condition (treated *versus* untreated) is indicated in parentheses. B: Functional categorization of the differentially expressed genes based on corresponding encoded (or putatively encoded) proteins. Only enriched categories (p-value < 0.05) are represented.

The functional categorization of the differentially expressed genes is represented in Fig. 3B. Genes with altered expression under fluconazole treatment were grouped into the single category “membrane lipid metabolism”. This category includes genes of the ergosterol biosynthesis pathway, such as *ERG11, ERG1, ERG2, CYB5*, and *NCP1*, which were upregulated (Table S3), as expected and reported by other authors (47, 48).

The genes differentially expressed after copper and Cu+Fluc treatment fall into nine categories (Fig. 3B). Both treatments showed significant enrichment in broad processes like metabolism, stress response, transcriptional control, and protein synthesis, consistent with a strong genetic reprogramming that cells should undergo. Included in the “homeostasis of metal ions” category are (i) genes involved in the copper-detoxification response, such as those encoding the metallothioneins (*MT-I, MT-IIA, MT-IIB*) and the transcriptional regulator of copper detoxification, *AMT1* (44, 45), and (ii) genes involved in copper transport and compartmentalization, such as *CTR1* and *CTR2*, which are homologs of the *S. cerevisiae* genes coding for the high-affinity copper importer and the vacuolar low-affinity copper transporter, respectively (49, 50). As anticipated, the first set of genes was upregulated, while the second was downregulated (Fig. 4). Under this category, we also find genes involved in Fe-S cluster biogenesis (*ISU1, ISA1, ISA2, CIA1, CIA2, NPB35*). Fe-S clusters are primary targets of copper toxicity (51) and their involvement in fluconazole tolerance has recently been demonstrated (27). Another group of genes comprised in this category code for proteins whose orthologs play a role in zinc homeostasis (52). These genes, which include *ZAP1*, the regulator of zinc homeostasis, and some of its putative target genes, were downregulated in copper and, more prominently, in Cu+Fluc conditions (Fig. 4), suggesting that zinc homeostasis might be especially affected by the latter treatment. Remarkably, *ZAP1* was also downregulated in cells exposed to fluconazole alone (Fig. 4).

**FIG 4.**
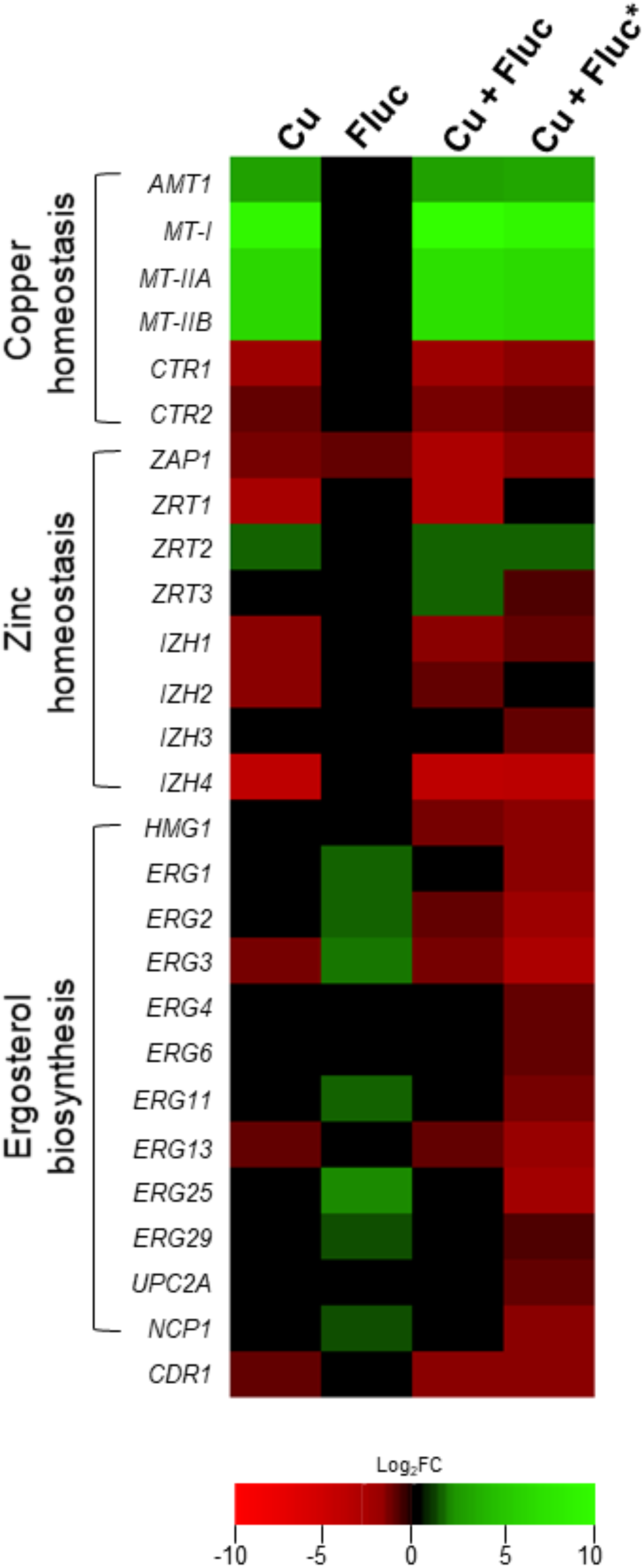
Heatmap depicting *Candida glabrata* genes belonging to the subcategories ergosterol biosynthesis, copper and zinc homeostasis, which were differentially expressed under Cu, Fluc or Cu+Fluc conditions. For each condition, the log_2_ fold change (FC) of the selected transcripts is indicated using a colour code (see the colour bar). Genes significantly up-(log_2_FC > 1) or downregulated (log_2_FC < -1) in response to Cu, Fluc or Cu+Fluc (treated *versus* untreated conditions) are shaded in green or red, respectively. Genes that were not differentially expressed between the treated and untreated conditions appear in black (−1 < log_2_FC < 1). The colours depicted in the Cu+Fluc* column indicate genes differentially expressed in Cu+Fluc conditions compared to Fluc conditions alone. *S. cerevisiae* gene names were used whenever the corresponding *C. glabrata* homolog name was not assigned.

The “membrane lipid metabolism” category was enriched under copper and Cu+Fluc treatment. Interestingly, we noticed that several genes of the ergosterol biosynthesis pathway were repressed in response to copper and Cu+Fluc (Fig. 4). This observation was even more striking when we compared gene expression under Cu+Fluc with that of cells only treated with fluconazole (Fig. 4 and Table S5). Clearly, the combination of copper and fluconazole led to the transcriptional repression of many genes relevant for ergosterol biosynthesis, among which were *UPC2A* and *ERG11* that encode the transcriptional regulator of the pathway (53) and the target of fluconazole (7), respectively.

The “stress response” category comprises several genes involved in oxidative stress detoxification (*CTA1, TRX2, GLR1, GSH2, TRR1*), found induced in response to both copper and Cu+Fluc (Tables S2 and S4), possibly to counteract the oxidative damage imposed by the metal (Fig. 2B). The gene encoding the ATP binding cassette (ABC) transporter Cdr1, which is linked with azole resistance in *Candida glabrata* (14), also appears under this category. Surprisingly, its expression was downregulated both by copper (log_2_FC_Cu_ -1.38) and Cu+Fluc (log_2_FC_Cu+Fluc_ -1.74), contrasting with its upregulation following treatment with fluconazole (14).

In summary, this analysis pinpoints two pathways known to be relevant for fluconazole adaptation that, according to the transcriptional data, might be negatively affected by the presence of copper or even worsen under Cu+Fluc conditions, namely (i) drug detoxification through Cdr1 and (ii) ergosterol biosynthesis. In addition, zinc homeostasis may be deregulated when copper and fluconazole are combined, which might pose a threat because zinc is an essential metal whose intracellular levels need to be delicately balanced.

### 3.4 Copper excess interferes with Pdr1 activity and leads to the accumulation of fluconazole

Confirming the RNA-Seq data, qRT-PCR analysis indicated that the transcript levels of *CDR1* were noticeably downregulated in response to copper and Cu+Fluc (Fig. 5A). We next tested whether the repression of *CDR1* expression by copper affected the accumulation of fluconazole. For that, we quantified the intracellular levels of fluconazole by HPLC. Consistent with *CDR1* repression, we observed that cells co-treated with copper accumulate higher levels of fluconazole (Fig. 5B).

**FIG 5.**
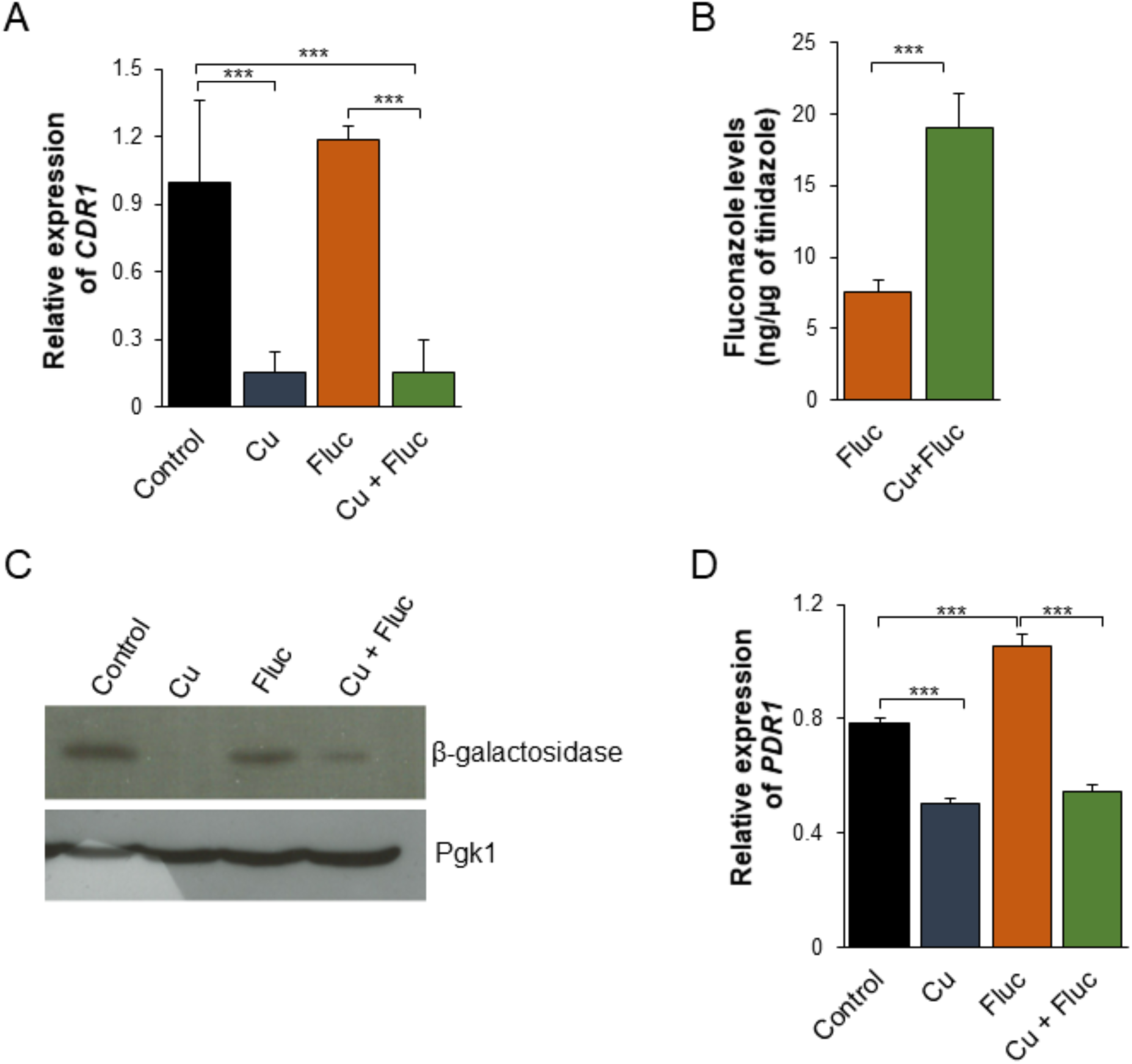
Copper treatment inhibits Pdr1 activity, downregulates *CDR1* expression and impairs fluconazole detoxification. A: The expression of the *CDR1* gene in cells untreated (Control) or treated with 1.25 mM CuSO_4_ (Cu), 32 µg/mL fluconazole (Fluc) or both (Cu+Fluc), was analyzed by qRT-PCR. B: The accumulation of fluconazole in cells untreated (Control) or treated with 625 µM of CuSO_4_ (Cu), 32 µg/mL fluconazole (Fluc) or both (Cu+Fluc) was measured by HPLC. The amount of fluconazole was normalized relatively to tinidazole (internal standard) C: *S. cerevisiae* EGY48 strain transformed with CgPdr1-*lexA* and pSH18-34 was grown until mid-log phase in selective media and treated (or left untreated) with 625 µM CuSO_4_ (Cu), 64 µg/mL fluconazole (Fluc) or both (Cu+Fluc) for 2 h. β-galactosidase protein levels were analyzed by Western blot. Pgk1 was used as loading control. D: The expression of the *PDR1* gene was assessed by qRT-PCR. Significance of differences was calculated using student’s T-test (*** p-value < 0.005).

In *Candida glabrata*, the Zn_2_Cys_6_ transcription factor Pdr1 regulates azole resistance through the induction of multidrug efflux genes, among which is *CDR1* (13, 46). Given the ability of copper to cause mismetallation and interfere with the activity of zinc-finger transcription factors (54), we decided to test whether the metal could be affecting the transactivation activity of Pdr1 under the conditions tested. To this end, we first generated a construct containing the *Candida glabrata PDR1* ORF fused to the *LexA* ORF regulated by the *ADH* promoter (CgPdr1-*LexA*). The *S. cerevisiae* EGY48 strain was then co-transformed with CgPdr1-*LexA* and the pSH18-34 plasmid, which contains the *LacZ* reporter gene, encoding β-galactosidase under the control of 8 L*exA* operators. The expression of the reporter gene, which reflects the ability of CgPdr1 to recruit the basal transcription machinery, was assessed by Western blotting (Fig. 5C). The results showed that copper, alone or combined with fluconazole, decreased the levels of β-galactosidase, suggesting that excessive levels of the metal (Fig. 2A) negatively affect the activity of Pdr1.

As the activity of Pdr1 is controlled by several factors, including transcriptional autoregulation (recently reviewed in (55)), we decided to measure the expression of *PDR1* by qRT-PCR under copper, fluconazole, Cu+Fluc and control conditions. Although *PDR1* was not considered differentially expressed in our RNA-Seq analysis, by qRT-PCR, we observed that, like *CDR1* (Fig. 5A), its expression was partly inhibited after treatment with copper, even in the presence of fluconazole (Fig. 5D).

Altogether, these results demonstrate that copper represses the expression of *CDR1*, which should explain the accumulation of the drug when copper and fluconazole are combined. The fact that the transactivation activity of Pdr1 is decreased under copper conditions further supports the idea that *CDR1* repression and fluconazole accumulation can be a consequence of Pdr1 inhibition imposed by excessive levels of the metal.

### 3.5 The combination of copper and fluconazole affects ergosterol metabolism

In response to ergosterol depletion caused by azoles, yeast cells increase the expression of the genes involved in the ergosterol biosynthesis pathway (*ERG* genes) (47, 48). Interestingly, our data shows that the combination of copper and fluconazole prevents this upregulation (Fig. 4). Accordingly, we found that *ERG11* mRNA levels were slightly but significantly downregulated in the presence of copper, and the combination of the metal with fluconazole precluded the upregulation of *ERG11*, observed when fluconazole was added to the cultures (Fig. 6A). Corroborating the transcriptional analyses, we found that Erg11 protein levels were compromised after co-treatment compared to fluconazole treatment alone (Fig. 6B). Contrary to our expectations, however, Cu+Fluc-treated cells possessed higher ergosterol levels than cells treated exclusively with fluconazole, as measured by HPLC (Fig. 6C).

**FIG 6.**
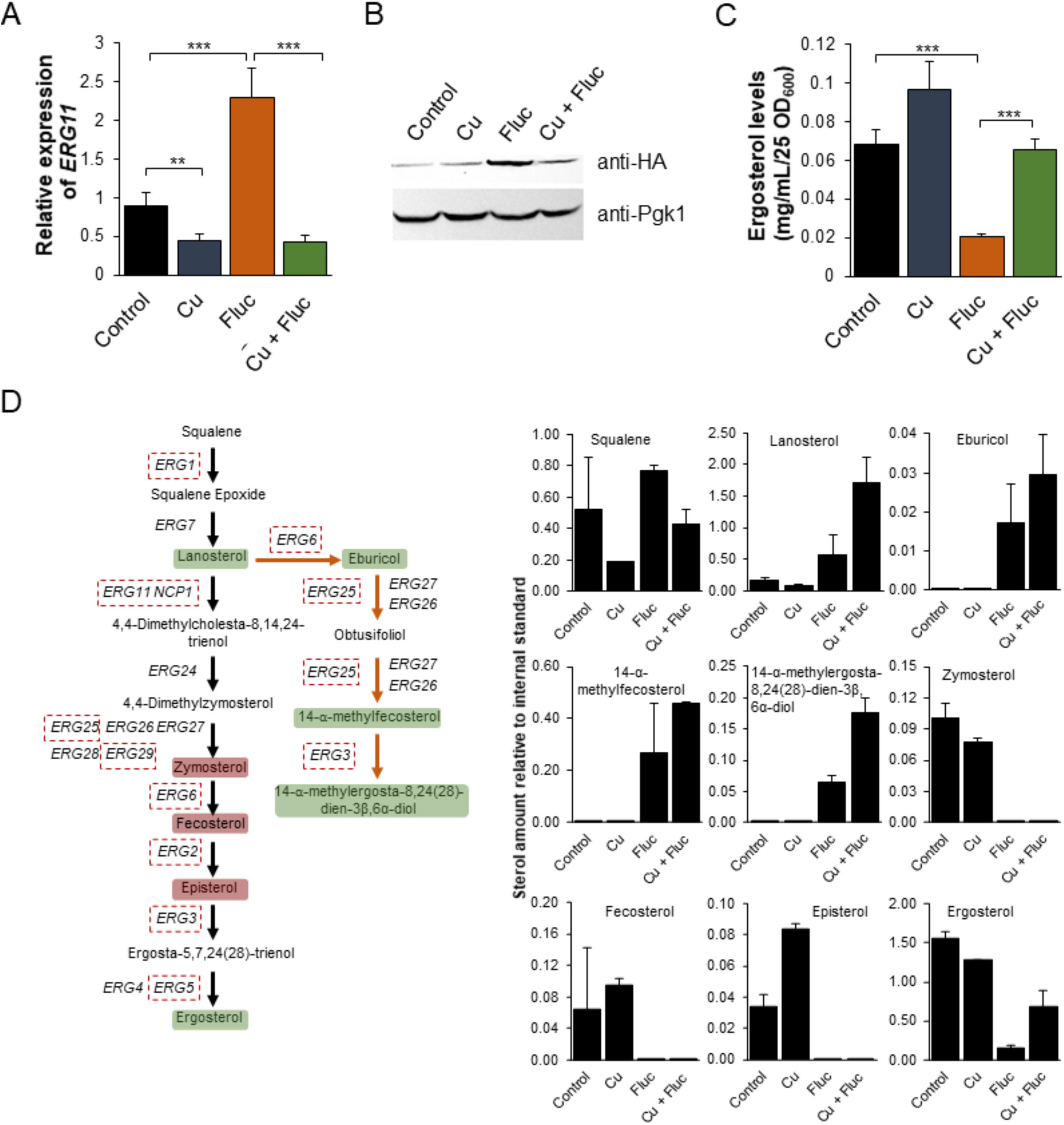
The combination of copper and fluconazole affects ergosterol metabolism. A: The expression of the *ERG11* gene was determined by qRT-PCR in *Candida glabrata* cells untreated (Control) or treated with 1.25 mM CuSO_4_ (Cu), 32 µg/mL fluconazole (Fluc) or with the two compounds (Cu+Fluc.). Western blot analyses of a HA-tagged version of Erg11, after treatment with 625 µM CuSO4, 32 µg/mL fluconazole or both for 2h 30 m. C: The ergosterol content of cells left untreated (Control) or treated with 625 µM of CuSO_4_, 32 µg/mL fluconazole (Fluc) or Cu+Fluc was measured by HPLC. Significance of differences was calculated using student’s T-test (**p-value<0.05; ***p-value<0.005). D: Schematic representation of the effect of Cu+Fluc on the biosynthesis of ergosterol. Dashed red boxes indicate the genes downregulated under Cu+Fluc treatment compared to fluconazole treatment, according to RNA-Seq analyses. Green boxes indicate sterol metabolites, which are more abundant after Cu+Fluc treatment, as compared to cells treated with fluconazole alone. Red boxes indicate sterol metabolites undetected by GC-MS under Cu+Fluc and Fluc conditions. The graphs indicate the relative amount of sterol intermediates in cells left untreated (Control) or treated with 625 µM of CuSO_4_ (Cu), 32 µg/mL fluconazole (Fluc) Cu+Fluc, as determined by GC-MS. Cholestane was used as an internal standard.

To have a comprehensive understanding of the impact of copper and fluconazole on ergosterol metabolism, we analyzed the sterol profile of *Candida glabrata* exposed to each condition, by GC-MS. A scheme summarizing the GC-MS results integrated with the transcriptomic data can be found in Fig. 6D. Lanosterol, the substrate of Erg11, as well as the intermediates of the ergosterol alternative pathway (eburicol, 14-α-methylfecosterol and 14-methylergosta-8,24(28)-dien-3β,6α-diol), which are known to compromise the function of the plasma membrane (7, 8), were not detected under copper and control conditions, but accumulated when cells were treated with fluconazole. The levels of these intermediates were overall higher under the conditions of Cu+Fluc than fluconazole alone (Fig. 6D), which is in line with the greater accumulation of fluconazole in the former condition (Fig. 5B). Zymosterol and fecosterol (late-stage intermediates downstream of lanosterol) could not be detected in any of the fluconazole conditions (Fig. 6D). Corroborating the HPLC analysis (Fig. 6C), by GC-MS, we confirmed that copper alleviates the ergosterol depletion caused by fluconazole (Fig. 6D).

As alterations in the sterol content of the yeast cell can affect membrane proteins that build and shape the cell wall and extracellular matrices (56), we used scanning electron microscopy to evaluate the impact of copper and fluconazole on cell surface morphology. *C. glabrata* untreated cells have a well-defined oval shape with a smooth surface (Fig. 7A), whereas copper- and fluconazole-treated cells sporadically exhibited a rough and irregular surface. In contrast, more than 50% of the cells treated with copper and fluconazole simultaneously have pronounced surface irregularities (Fig. 7A). The morphology of untreated and treated cells was also observed using transmission electron microscopy (Fig. 7). Untreated cells have normal cell walls (cw) with a thin electron-dense outer layer with protruding fibrillar structures (fs). A continuous inner plasma membrane (pm) delimits an electron-dense and conserved cytoplasm in which nucleus (n) and vacuoles (v) are visible. Treatment with copper and fluconazole induced structural changes at the periphery and inside the cells, which were more pronounced when both treatments were combined. Among the ultrastructural alterations were: irregular cell wall and plasma membrane (all treatments), decreased cytoplasmic density and disorganization of the cytoplasm (all treatments), clear defects in the cell wall (Cu and Cu+Fluc treatments, black arrows in Fig. 7) and, in some cells, loss of cell wall integrity (Cu+Fluc treatment, red arrows in Fig. 7). Both the accumulation of copper (Fig. 2A) and the alteration of sterol metabolism, despite the higher levels of ergosterol (Fig. 6D), should certainly contribute to the Cu+Fluc phenotype.

**FIG 7.**
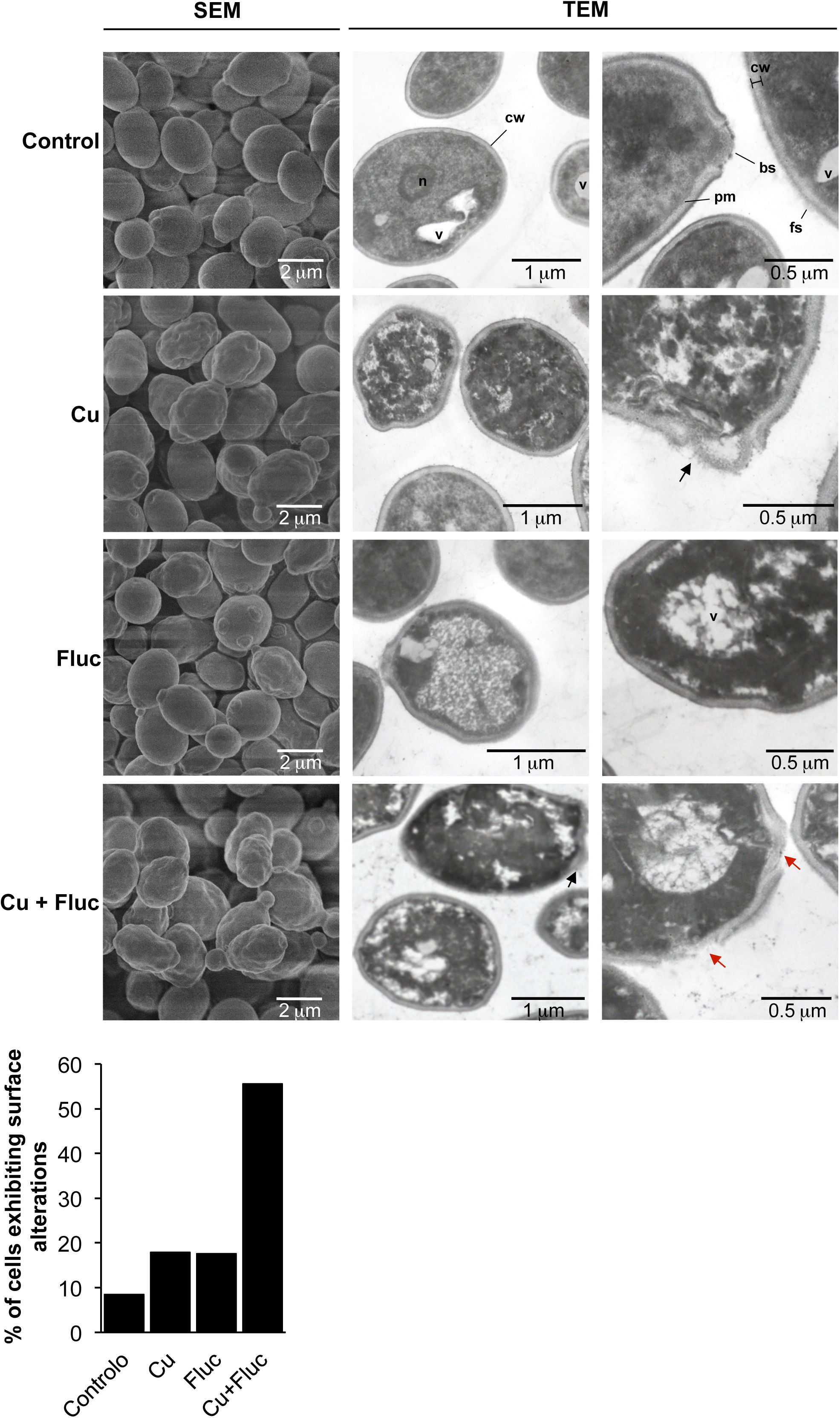
Scanning electron microscopy (SEM) and transmission electron microscopy (TEM) images of *Candida glabrata* cells left untreated (Control), or treated with 625 µM CuSO_4_ (Cu), 32 µg/mL fluconazole (Fluc) or both (Cu+Fluc) for 24 h. Control cells have regular cell walls (cw), from which fibrillar structures (f) protrude, well defined and continuous plasma membrane (pm) and visible vacuoles (v), nucleus (n) and bud scars (bs). Cells treated with Cu, Fluc or Cu+Fluc exhibit irregular surfaces and disorganized cytoplasm. Cu and Cu+Fluc stresses also induce defects in the cell wall (black arrows) and in the latter, loss of cell wall integrity (red arrows) was observable in some cells. The graph indicates the percentage of cells exhibiting surface abnormalities. A total of 120 cells per condition were imaged by SEM and those with irregular surfaces were counted.

### 3.6 Co-treatment of *Candida glabrata* cells with copper and fluconazole causes zinc depletion

The striking observation that several genes presumably involved in zinc sensing and transport were downregulated after fluconazole, copper and Cu+Fluc treatment (Fig. 4) led us to investigate whether zinc homeostasis could be affected. Included in this set of genes was *ZAP1* that, in yeasts, encodes a zinc-finger transcription factor that acts as master regulator of zinc homeostasis (57).

After confirming the transcriptomic data, by qRT-PCR (Fig. 8A), we further explored the relevance of Zap1, by evaluating how *ZAP1* deletion affected yeast survival in each individual or combined condition. We tested the knockout effect of the gene in *Candida glabrata* and in *Saccharomyces cerevisiae*, two phylogenetically closely related species (58). Reinforcing the role of Zap1 in yeast survival under these stress conditions, we found that *ScΔzap1* cells were slightly sensitive to fluconazole and strongly sensitive to copper and Cu+Fluc, with the latter triggering a more pronounced effect (Fig. 8B). Mutant *CgΔzap1* was also sensitive to all stresses, but more prominently to fluconazole and Cu+Fluc (Fig. 8C). The tolerance to fluconazole and Cu+Fluc upon reintroduction of Cg*ZAP1* was restored to that of the parent strain (Fig.8C).

**FIG 8.**
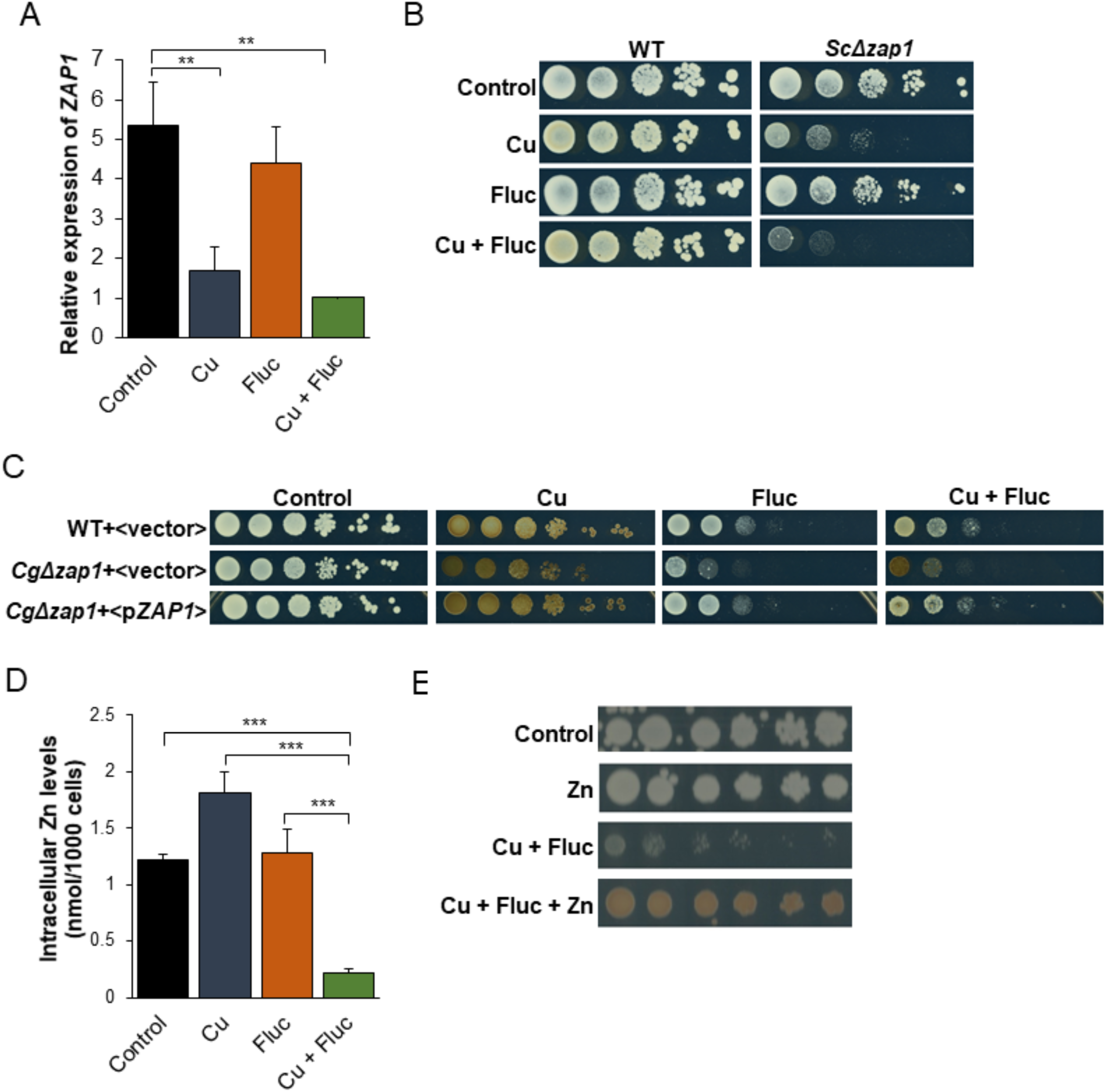
Zinc homeostasis and the transcription factor Zap1 are relevant for cells to thrive under Cu+Fluc conditions. A: The expression of *ZAP1* gene in *Candida glabrata* cells left untreated (Control), treated with 1.25 mM CuSO_4_ (Cu), 32 µg/mL fluconazole (Fluc) or with a combination of Cu and Fluc (Cu+Fluc) was evaluated by qRT-PCR. B: Growth of *Saccharomyces cerevisiae* wild-type (WT:BY4742) and Δ*zap1* mutant strains in SC agar plates (Control) containing 4 µg/mL of fluconazole (Fluc), 625 µM CuSO_4_ (Cu) or a combination of both (Cu+Fluc). Growth was recorded after 96 h at 30 ºC. C: Growth of *Candida glabrata* wild-type (WT) and Δ*zap1* mutant strains transformed with a plasmid containing the Cg*ZAP1* gene under the control of its native promoter (<p*ZAP1*>) or with the empty vector (<vector>) in SC agar plates lacking tryptophan (Control) containing 32 µg/mL of fluconazole (Fluc), 625 µM CuSO_4_ (Cu) or a combination of both (Cu+Fluc), after 48 h at 37 ºC. D: The zinc content of cells untreated (Control) or treated with 625 µM CuSO_4_ (Cu), 32 µg/mL fluconazole (Fluc) or a combination of both was measured by ICP-AES. E: Effect of zinc on the growth sensitivity of *Candida glabrata* cells to Cu+Fluc. Cells were serial diluted and spotted on SD plates containing a mixture of fluconazole (32 µg/mL) and CuSO_4_ (625 µM), supplemented (Cu+Fluc+Zn) or not (Cu+Fluc) with 4 mM of ZnSO_4_. Growth was recorded after 48 h at 37ºC.

We next assessed by ICP-AES whether the downregulation of Cg*ZAP1* (Fig. 8D) could impact zinc accumulation. As shown in Fig. 8D, the levels of zinc were found to be strongly depleted in *C. glabrata* cells treated with Cu+Fluc but unchanged or even increased under fluconazole and copper conditions, respectively. Zinc supplementation rescued the growth of *Candida glabrata* cells treated with Cu+Fluc and increased the brownish pigmentation associated with CuS mineralization (Fig. 8E).

Together, these results show that cells exposed to Cu+Fluc fail to uptake zinc and are strongly deprived of this metal, suggesting that Zap1 (and zinc) is important for an adequate response to the stress imposed by the combination of copper and fluconazole.

## 4. Discussion

In recent years, there was a growing interest in exploiting the antifungal properties of copper. Trojan horse strategies that deceive the fungus copper homeostatic mechanisms (20, 29) or the combination of copper with available antifungals (18, 19, 59) have proved to be effective against several fungal species. In particular, copper has been shown to potentiate the activity of fluconazole (18, 19, 43), one of the most prescribed antifungals worldwide. However, despite its interest and great promise, the mechanism underlying this effect remains unknown.

In this work, we have investigated the molecular basis that supports the synergistic effect observed between copper and fluconazole in *Candida glabrata* (Fig. 1), a yeast intrinsically tolerant to fluconazole and frequently associated with invasive candidiasis (5, 9, 10).

The analyses of the transcriptome of *Candida glabrata* (Figs. 3 and 4) revealed that the gene encoding a multidrug ABC transporter essential for fluconazole detoxification, *CDR1* (14), is repressed in response to copper, even in the presence of fluconazole (Fig. 5A and Tables S2 and S4). As copper impairs the activity of Pdr1 (Fig. 5C), a Zn_2_Cys_6_ transcription factor known to regulate *CDR1* expression (13, 46), it is possible that *CDR1* repression is a consequence of copper-mediated Pdr1 inhibition. Copper can displace zinc from zinc-finger motifs of transcription factors because of its strong affinity for cysteine-rich domains, which causes structural alterations that lead to the loss of the DNA binding activity (54). Although in this study we have not addressed the effect of copper on Pdr1 binding to DNA, it is clear that the metal also affects the transactivation potential of Pdr1 (Fig. 5C), that is, Pdr1’s ability to recruit the basal transcriptional machinery. As a result, in the presence of copper and fluconazole, *Candida glabrata* fails to properly activate the fluconazole detoxification response and, therefore, accumulates higher amounts of the drug (Fig. 5B).

In agreement with the greater intracellular content of fluconazole, we found that Cu+Fluc treatment elicits a higher accumulation of lanosterol and toxic methylated sterol intermediates (eburicol and 14-α-methylfecosterol). These sterol metabolites alter the physicochemical properties of the plasma membrane, thus affecting membrane-associated processes like transport, growth, and division (7, 8). Unexpectedly, under co-treatment conditions we found that fluconazole no longer causes ergosterol depletion, as measured by HPLC and confirmed by GC-MS (Fig. 6). Although we cannot explain this observation at present, we cannot rule out the possibility of the existence of an as-yet unveiled alternative pathway, specifically activated by copper, which enables the conversion of sterol intermediates into ergosterol, as previously proposed by other authors for other fungi (60, 61).

Further corroborating the strong deregulation of the sterol metabolism imposed by Cu+Fluc, we noticed that many genes of the ergosterol pathway were downregulated as opposed to fluconazole treatment (Figs. 4 and 6). In *Candida glabrata*, sterol metabolism is controlled by two Zn_2_Cys_6_ transcription factors, Upc2a and Upc2b, orthologs of *S. cerevisiae* Upc2 and Emc22, respectively (62). Upc2a is essential for the activation of *ERG* genes in response to fluconazole treatment, while Upc2b is involved mainly in the uptake of exogenous sterols (62). Studies in *S. cerevisiae* revealed that Upc2 regulates its own expression (63) and that, in response to high levels of ergosterol, it translocates to the cytoplasm (64). In this sense, it is possible that under Cu+Fluc treatment, the higher levels of ergosterol (Fig. 6) block the activation of Upc2a that would, under normal fluconazole conditions, activate the *ERG* genes (47). Another possibility is that copper interferes with the activity of Upc2a *via* zinc-finger disruption. Both options would also explain why the expression of *UPC2A* is downregulated by copper in the presence of fluconazole (Fig. 4).

The accumulation of toxic sterol intermediates (65), together with the ability of copper to affect membrane permeability and fluidity because of lipid peroxidation and interaction with phospholipids, is highly indicative that membrane alterations might be central for the synergistic effect between copper and fluconazole. Supporting this idea, SEM and TEM analysis revealed pronounced cell surface irregularities, which were more accentuated under Cu+Fluc conditions (Fig. 7). As the activity of membrane transporters depends on their correct insertion into the membrane bilayer and their specific interactions with lipids (66), it is possible that Cu+Fluc co-treatment compromises metal transport activity, which would explain the decreased accumulation of copper observed in cells treated with Cu+Fluc (Fig. 2A). The loss of pigmentation of Cu+Fluc treated cells (Fig. 1A) may also be a consequence of impaired function of cell surface-bound enzymes that mineralize CuS to confer resistance to this metal (42).

Zap1 is the master regulator of zinc homeostasis in yeasts (52, 67), and here we showed that it is important for cells to deal with copper and fluconazole toxicity. The treatment with copper, fluconazole, or a combination of both, represses the expression of *ZAP1*, with such effect being more prominent under copper and Cu+Fluc conditions (Figs. 4 and 8). Zap1 contains seven zinc-finger motifs in its structure, two of which acting as zinc sensors (68) and, like Pdr1 (69), it regulates its own expression (70).Therefore, Zap1 activity might as well be impaired by the excess of copper, which would explain the strong *ZAP1* repression under copper and Cu+Fluc conditions (Fig. 8A). However, despite a strong downregulation of *ZAP1* by copper, zinc depletion was only evident when cells were simultaneously treated with fluconazole (Cu+Fluc), but not with the metal alone (Fig. 8D). We reason that the possible malfunction of membrane metal transporters, due to exacerbated membrane defects observed under Cu+Fluc (Fig. 7), may justify this observation. The requirement of zinc for an adequate response to Cu+Fluc is reinforced by the observation that zinc supplementation restores growth under these conditions (Fig. 8E). Because zinc is a ubiquitous element, essential for the function of many transcription factors, including Upc2a/b and Pdr1, we propose that, to cope with Cu+Fluc treatment, *C. glabrata* requires adequate zinc levels for these transcription factors to function properly.

A recent study by Hunsaker *et. al* evaluated transcriptomic alterations of *Candida albicans* cells exposed to a combination of copper and fluconazole (71). Unlike *Candida glabrata*, under these conditions, *C. albicans* cells do not reduce the expression of genes involved in drug detoxification (*CDR1* and *TAC1*), ergosterol metabolism (*ERG* genes) or zinc homeostasis. Also contrasting with our data (Fig. 2), in a previous study, the authors showed that co-treatment increases copper accumulation when compared to copper treatment alone (43). Therefore, it appears that the mechanisms underlying the synergistic effect between copper and fluconazole are quite distinct in *C. glabrata* and *C. albicans*.

In conclusion, our work suggests that copper may negatively impact on the activity of several zinc-finger transcription factors that control important pathways for fluconazole detoxification: azole efflux (Pdr1), ergosterol biosynthesis (Upc2), and zinc homeostasis (Zap1). Therefore, when copper and fluconazole are combined, cells fail to detoxify the drug and accumulate toxic intermediates of the sterol metabolism. This accumulation, possibly together with copper-induced lipid damages, should affect the activity of membrane metal transporters. Zinc transporters malfunction together with the downregulation of *ZAP1* and its target genes should impede zinc uptake, leading to zinc depletion, which further sensitizes *Candida glabrata* cells to copper and fluconazole co-treatment.

Overall, our findings strongly support continued investment in bi-functional drugs, combining azole and copper chelating groups. Such a strategy, while ensuring a synchronized uptake of both compounds, would deceive the yeast copper homeostatic mechanisms, and possibly decrease the effective copper concentration needed to achieve the same synergistic effect, which would certainly reduce the toxicity to human cells.

## 5. Acknowledgments

We thank Cristina Leitão (Research Facilities ITQB NOVA) for technical assistance in HPLC. The authors are indebted to Professor Claudina Rodrigues-Pousada for her guidance and support throughout the years as the leader of the Genomics and Stress laboratory at ITQB-NOVA.The authors acknowledge the assistance of Beatriz Carmo in the construction of the *CgΔzap1* mutant. We are grateful to Professor Scott Moye-Rowley (University of Iowa) and Professor Patrick Van Dijck (KU Leuven) for providing the BVGC3 and Cg*Δcdr1* strains, respectively.

## 7. Supporting information

This article contains supporting information.

## References

1. Bongomin F, Gago S, Oladele RO, Denning DW. 2017. Global and Multi-National Prevalence of Fungal Diseases-Estimate Precision. J Fungi (Basel) 3.

2. Pfaller MA, Diekema DJ. 2007. Epidemiology of invasive candidiasis: a persistent public health problem. Clin Microbiol Rev 20:133–63.

3. Paiva JA, Pereira JM, Tabah A, Mikstacki A, de Carvalho FB, Koulenti D, Ruckly S, Cakar N, Misset B, Dimopoulos G, Antonelli M, Rello J, Ma X, Tamowicz B, Timsit JF. 2016. Characteristics and risk factors for 28-day mortality of hospital acquired fungemias in ICUs: data from the EUROBACT study. Crit Care 20:53.

4. Tabah A, Koulenti D, Laupland K, Misset B, Valles J, Bruzzi de Carvalho F, Paiva JA, Cakar N, Ma X, Eggimann P, Antonelli M, Bonten MJ, Csomos A, Krueger WA, Mikstacki A, Lipman J, Depuydt P, Vesin A, Garrouste-Orgeas M, Zahar JR, Blot S, Carlet J, Brun-Buisson C, Martin C, Rello J, Dimopoulos G, Timsit JF. 2012. Characteristics and determinants of outcome of hospital-acquired bloodstream infections in intensive care units: the EUROBACT International Cohort Study. Intensive Care Med 38:1930–45.

5. Pfaller MA, Andes DR, Diekema DJ, Horn DL, Reboli AC, Rotstein C, Franks B, Azie NE. 2014. Epidemiology and outcomes of invasive candidiasis due to non-albicans species of Candida in 2,496 patients: data from the Prospective Antifungal Therapy (PATH) registry 2004-2008. PLoS One 9:e101510.

6. Pappas PG, Kauffman CA, Andes DR, Clancy CJ, Marr KA, Ostrosky-Zeichner L, Reboli AC, Schuster MG, Vazquez JA, Walsh TJ, Zaoutis TE, Sobel JD. 2016. Clinical Practice Guideline for the Management of Candidiasis: 2016 Update by the Infectious Diseases Society of America. Clin Infect Dis 62:e1–50.

7. Kelly SL, Lamb DC, Corran AJ, Baldwin BC, Kelly DE. 1995. Mode of action and resistance to azole antifungals associated with the formation of 14 alpha-methylergosta-8,24(28)-dien-3 beta,6 alpha-diol. Biochem Biophys Res Commun 207:910–5.

8. Watson PF, Rose ME, Ellis SW, England H, Kelly SL. 1989. Defective sterol C5-6 desaturation and azole resistance: a new hypothesis for the mode of action of azole antifungals. Biochem Biophys Res Commun 164:1170–5.

9. Whaley SG, Rogers PD. 2016. Azole Resistance in Candida glabrata. Curr Infect Dis Rep 18:41.

10. Pfaller MA, Diekema DJ, Turnidge JD, Castanheira M, Jones RN. 2019. Twenty Years of the SENTRY Antifungal Surveillance Program: Results for. Open Forum Infect Dis 6:S79–S94.

11. Kasper L, Seider K, Hube B. 2015. Intracellular survival of Candida glabrata in macrophages: immune evasion and persistence. FEMS Yeast Res 15:fov042.

12. Tsai HF, Krol AA, Sarti KE, Bennett JE. 2006. Candida glabrata PDR1, a transcriptional regulator of a pleiotropic drug resistance network, mediates azole resistance in clinical isolates and petite mutants. Antimicrob Agents Chemother 50:1384–92.

13. Vermitsky JP, Earhart KD, Smith WL, Homayouni R, Edlind TD, Rogers PD. 2006. Pdr1 regulates multidrug resistance in Candida glabrata: gene disruption and genome-wide expression studies. Mol Microbiol 61:704–22.

14. Sanglard D, Ischer F, Calabrese D, Majcherczyk PA, Bille J. 1999. The ATP binding cassette transporter gene CgCDR1 from Candida glabrata is involved in the resistance of clinical isolates to azole antifungal agents. Antimicrob Agents Chemother 43:2753–65.

15. Smith AD, Logeman BL, Thiele DJ. 2017. Copper Acquisition and Utilization in Fungi. Annu Rev Microbiol 71:597–623.

16. García-Santamarina S, Thiele DJ. 2015. Copper at the Fungal Pathogen-Host Axis. J Biol Chem 290:18945–53.

17. Mackie J, Szabo EK, Urgast DS, Ballou ER, Childers DS, MacCallum DM, Feldmann J, Brown AJ. 2016. Host-Imposed Copper Poisoning Impacts Fungal Micronutrient Acquisition during Systemic Candida albicans Infections. PLoS One 11:e0158683.

18. Ząbek A, Nagaj J, Grabowiecka A, Dworniczek E, Nawrot U, Młynarz P, Jeżowska-Bojczuk M. 2015. Activity of fluconazole and its Cu(II) complex towards Candida species. Medicinal Chemistry Research 24:2005–2010.

19. Hunsaker EW, Franz KJ. 2019. Copper potentiates azole antifungal activity in a way that does not involve complex formation. Dalton Trans 48:9654–9662.

20. Hunsaker EW, McAuliffe KJ, Franz KJ. 2020. Fluconazole analogues with metal-binding motifs impact metal-dependent processes and demonstrate antifungal activity in Candida albicans. J Biol Inorg Chem.

21. Menezes RA, Amaral C, Delaunay A, Toledano M, Rodrigues-Pousada C. 2004. Yap8p activation in Saccharomyces cerevisiae under arsenic conditions. FEBS Lett 566:141–6.

22. Pimentel C, Vicente C, Menezes RA, Caetano S, Carreto L, Rodrigues-Pousada C. 2012. The role of the Yap5 transcription factor in remodeling gene expression in response to Fe bioavailability. PLoS One 7:e37434.

23. Collart MA, Oliviero S. 1993. Preparation of Yeast RNA. Current Protocols in Molecular Biology 23:13.12.1-13.12.5.

24. da Silva SM, Batista-Nascimento L, Gaspar-Cordeiro A, Vernis L, Pimentel C, Rodrigues-Pousada C. 2018. Transcriptional regulation of FeS biogenesis genes: A possible shield against arsenate toxicity activated by Yap1. Biochim Biophys Acta Gen Subj 1862:2152–2161.

25. Gaspar-Cordeiro A, Marques Caetano S, Amaral C, Rodrigues-Pousada C, Pimentel C. 2018. Ace1 prevents intracellular copper accumulation by regulating Fet3 expression and thereby restricting Aft1 activity. FEBS J 285:1861–1872.

26. Morio F, Pagniez F, Lacroix C, Miegeville M, Le Pape P. 2012. Amino acid substitutions in the Candida albicans sterol Δ5,6-desaturase (Erg3p) confer azole resistance: characterization of two novel mutants with impaired virulence. J Antimicrob Chemother 67:2131–8.

27. Demuyser L, Swinnen E, Fiori A, Herrera-Malaver B, Vestrepen K, Van Dijck P. 2017. Mitochondrial Cochaperone Mge1 Is Involved in Regulating Susceptibility to Fluconazole in. mBio 8.

28. Müller C, Binder U, Bracher F, Giera M. 2017. Antifungal drug testing by combining minimal inhibitory concentration testing with target identification by gas chromatography-mass spectrometry. Nat Protoc 12:947–963.

29. Gaspar-Cordeiro A, da Silva S, Aguiar M, Rodrigues-Pousada C, Haas H, Lima LMP, Pimentel C. 2020. A copper(II)-binding triazole derivative with ionophore properties is active against Candida spp. J Biol Inorg Chem.

30. Pais P, Califórnia R, Galocha M, Viana R, Ola M, Cavalheiro M, Takahashi-Nakaguchi A, Chibana H, Butler G, Teixeira MC. 2020. Candida glabrata Transcription Factor Rpn4 Mediates Fluconazole Resistance through Regulation of Ergosterol Biosynthesis and Plasma Membrane Permeability. Antimicrob Agents Chemother 64.

31. Vu BG, Thomas GH, Moye-Rowley WSCP. 2019. Evidence that Ergosterol Biosynthesis Modulates Activity of the Pdr1 Transcription Factor in Candida glabrata. mBio 10.

32. Picelli S, Faridani OR, Björklund AK, Winberg G, Sagasser S, Sandberg R. 2014. Full-length RNA-seq from single cells using Smart-seq2. Nat Protoc 9:171–81.

33. Langmead B, Salzberg SL. 2012. Fast gapped-read alignment with Bowtie 2. Nat Methods 9:357–9.

34. Li H, Handsaker B, Wysoker A, Fennell T, Ruan J, Homer N, Marth G, Abecasis G, Durbin R, Subgroup GPDP. 2009. The Sequence Alignment/Map format and SAMtools. Bioinformatics 25:2078–9.

35. Liao Y, Smyth GK, Shi W. 2014. featureCounts: an efficient general purpose program for assigning sequence reads to genomic features. Bioinformatics 30:923–30.

36. Robinson MD, McCarthy DJ, Smyth GK. 2010. edgeR: a Bioconductor package for differential expression analysis of digital gene expression data. Bioinformatics 26:139–40.

37. Priebe S, Linde J, Albrecht D, Guthke R, Brakhage AA. 2011. FungiFun: a web-based application for functional categorization of fungal genes and proteins. Fungal Genet Biol 48:353–8.

38. Institute CLSICaLS. 2017. Reference method for broth dilution antifungal susceptibility testing of yeasts CLSI standard M27, 4th ed.

39. Fiori A, Van Dijck P. 2012. Potent synergistic effect of doxycycline with fluconazole against Candida albicans is mediated by interference with iron homeostasis. Antimicrob Agents Chemother 56:3785–96.

40. Marchetti O, Moreillon P, Glauser MP, Bille J, Sanglard D. 2000. Potent synergism of the combination of fluconazole and cyclosporine in Candida albicans. Antimicrob Agents Chemother 44:2373–81.

41. Lachke SA, Srikantha T, Tsai LK, Daniels K, Soll DR. 2000. Phenotypic switching in Candida glabrata involves phase-specific regulation of the metallothionein gene MT-II and the newly discovered hemolysin gene HLP. Infect Immun 68:884–95.

42. Yu W, Farrell RA, Stillman DJ, Winge DR. 1996. Identification of SLF1 as a new copper homeostasis gene involved in copper sulfide mineralization in Saccharomyces cerevisiae. Mol Cell Biol 16:2464–72.

43. Hunsaker EW, Franz KJ. 2019. Candida albicans reprioritizes metal handling during fluconazole stress. Metallomics 11:2020–2032.

44. Mehra RK, Thorvaldsen JL, Macreadie IG, Winge DR. 1992. Disruption analysis of metallothionein-encoding genes in Candida glabrata. Gene 114:75–80.

45. Zhou PB, Thiele DJ. 1991. Isolation of a metal-activated transcription factor gene from Candida glabrata by complementation in Saccharomyces cerevisiae. Proc Natl Acad Sci U S A 88:6112–6.

46. Vermitsky JP, Edlind TD. 2004. Azole resistance in Candida glabrata: coordinate upregulation of multidrug transporters and evidence for a Pdr1-like transcription factor. Antimicrob Agents Chemother 48:3773–81.

47. Henry KW, Nickels JT, Edlind TD. 2000. Upregulation of ERG genes in Candida species by azoles and other sterol biosynthesis inhibitors. Antimicrob Agents Chemother 44:2693–700.

48. Whaley SG, Caudle KE, Vermitsky JP, Chadwick SG, Toner G, Barker KS, Gygax SE, Rogers PD. 2014. UPC2A is required for high-level azole antifungal resistance in Candida glabrata. Antimicrob Agents Chemother 58:4543–54.

49. Dancis A, Haile D, Yuan DS, Klausner RD. 1994. The Saccharomyces cerevisiae copper transport protein (Ctr1p). Biochemical characterization, regulation by copper, and physiologic role in copper uptake. J Biol Chem 269:25660–7.

50. Rees EM, Lee J, Thiele DJ. 2004. Mobilization of intracellular copper stores by the ctr2 vacuolar copper transporter. J Biol Chem 279:54221–9.

51. Macomber L, Imlay JA. 2009. The iron-sulfur clusters of dehydratases are primary intracellular targets of copper toxicity. Proc Natl Acad Sci U S A 106:8344–9.

52. Zhao H, Butler E, Rodgers J, Spizzo T, Duesterhoeft S, Eide D. 1998. Regulation of zinc homeostasis in yeast by binding of the ZAP1 transcriptional activator to zinc-responsive promoter elements. J Biol Chem 273:28713–20.

53. Whaley SG, Caudle KE, Vermitsky JP, Chadwick SG, Toner G, Barker KS, Gygax SE, Rogers PDCP. 2014. UPC2A is required for high-level azole antifungal resistance in Candida glabrata. Antimicrob Agents Chemother 58:4543–54.

54. Predki PF, Sarkar B. 1994. Metal replacement in “zinc finger” and its effect on DNA binding. Environ Health Perspect 102 Suppl 3:195–8.

55. Moye-Rowley WS. 2019. Multiple interfaces control activity of the Candida glabrata Pdr1 transcription factor mediating azole drug resistance. Curr Genet 65:103–108.

56. Kodedová M, Sychrová H. 2015. Changes in the Sterol Composition of the Plasma Membrane Affect Membrane Potential, Salt Tolerance and the Activity of Multidrug Resistance Pumps in Saccharomyces cerevisiae. PLoS One 10:e0139306.

57. Eide DJ. 2009. Homeostatic and adaptive responses to zinc deficiency in Saccharomyces cerevisiae. J Biol Chem 284:18565–9.

58. Marcet-Houben M, Gabaldón T. 2009. The tree versus the forest: the fungal tree of life and the topological diversity within the yeast phylome. PLoS One 4:e4357.

59. Nagaj J, Starosta R, Szczepanik W, Barys M, Młynarz P, Jeżowska-Bojczuk M. 2012. The Cu(II)-fluconazole complex revisited. Part I: Structural characteristics of the system. J Inorg Biochem 106:23–31.

60. Fowler DM, Cooper SJ, Stephany JJ, Hendon N, Nelson S, Fields S. 2011. Suppression of statin effectiveness by copper and zinc in yeast and human cells. Mol Biosyst 7:533–44.

61. Huster D, Purnat TD, Burkhead JL, Ralle M, Fiehn O, Stuckert F, Olson NE, Teupser D, Lutsenko S. 2007. High copper selectively alters lipid metabolism and cell cycle machinery in the mouse model of Wilson disease. J Biol Chem 282:8343–55.

62. Nagi M, Nakayama H, Tanabe K, Bard M, Aoyama T, Okano M, Higashi S, Ueno K, Chibana H, Niimi M, Yamagoe S, Umeyama T, Kajiwara S, Ohno H, Miyazaki Y. 2011. Transcription factors CgUPC2A and CgUPC2B regulate ergosterol biosynthetic genes in Candida glabrata. Genes Cells 16:80–9.

63. Abramova NE, Cohen BD, Sertil O, Kapoor R, Davies KJ, Lowry CV. 2001. Regulatory mechanisms controlling expression of the DAN/TIR mannoprotein genes during anaerobic remodeling of the cell wall in Saccharomyces cerevisiae. Genetics 157:1169–77.

64. Yang H, Tong J, Lee CW, Ha S, Eom SH, Im YJ. 2015. Structural mechanism of ergosterol regulation by fungal sterol transcription factor Upc2. Nat Commun 6:6129.

65. Abe F, Usui K, Hiraki T. 2009. Fluconazole modulates membrane rigidity, heterogeneity, and water penetration into the plasma membrane in Saccharomyces cerevisiae. Biochemistry 48:8494–504.

66. Lv QZ, Yan L, Jiang YY. 2016. The synthesis, regulation, and functions of sterols in Candida albicans: Well-known but still lots to learn. Virulence 7:649–59.

67. Nobile CJ, Nett JE, Hernday AD, Homann OR, Deneault JS, Nantel A, Andes DR, Johnson AD, Mitchell AP. 2009. Biofilm matrix regulation by Candida albicans Zap1. PLoS Biol 7:e1000133.

68. Bird AJ, McCall K, Kramer M, Blankman E, Winge DR, Eide DJ. 2003. Zinc fingers can act as Zn2+ sensors to regulate transcriptional activation domain function. EMBO J 22:5137–46.

69. Khakhina S, Simonicova L, Moye-Rowley WS. 2018. Positive autoregulation and repression of transactivation are key regulatory features of the Candida glabrata Pdr1 transcription factor. Mol Microbiol 107:747–764.

70. Zhao H, Eide DJ. 1997. Zap1p, a metalloregulatory protein involved in zinc-responsive transcriptional regulation in Saccharomyces cerevisiae. Mol Cell Biol 17:5044–52.

71. Hunsaker EW, Yu CA, Franz KJ. 2021. Copper Availability Influences the Transcriptomic Response of Candida albicans to Fluconazole Stress. G3 (Bethesda) 11.

