## Supplementary material for "Copper Acts Synergistically with Fluconazole in *Candida glabrata* by Compromising Drug Efflux, Sterol Metabolism, and Zinc Homeostasis": Fig S1

Running title: Copper and fluconazole synergy in *Candida glabrata*

[1]Gaspar-Cordeiro, A., [1]Amaral, C., [1]Pobre, V., [1,2]Antunes, W., [1]Petronilho, A. [3]Paixão, P., [4]Alves de Matos, A.P., C., [1*]Pimentel, C.

[1] Instituto de Tecnologia Química e Biológica António Xavier, Universidade Nova de Lisboa, Av. República, 2780-157 Oeiras, Portugal.
[2] Centro de Investigação da Academia Militar (CINAMIL), Unidade Militar Laboratorial de Defesa Biológica e Química (UMLDBQ), Av. Dr. Alfredo Bensaúde, 1849-012 Lisboa, Portugal.
[3] Unidade de Infeção. Chronic Diseases Research Centre - CEDOC. NOVA Medical School/Faculdade de Ciências Médicas. Universidade NOVA de Lisboa. Lisboa; Laboratório de Patologia Clínica - SYNLAB. Hospital da Luz. Lisboa. Portugal.
[4] Egas Moniz Interdisciplinary Research Centre, Egas Moniz Higher Education Cooperative, Caparica, Portugal.

### Supporting Information Content:

- Table S1 – Strains used in this study.
- Table S2 - Genes differentially expressed in *Candida glabrata* cells treated with copper (submitted as .xslx)
- Table S3 - Genes differentially expressed in *Candida glabrata* cells treated with fluconazole (submitted as .xslx).
- Table S4 - Genes differentially expressed in *Candida glabrata* cells treated with copper and fluconazole (submitted as .xslx).
- Table S5 - Genes differentially expressed in *Candida glabrata* cells treated with copper and fluconazole in comparison to fluconazole alone (submitted as .xslx).
- Figure S1 - Effect of the combination of copper and fluconazole on the growth of A: different laboratory (HTL and HTU) and B: clinical strains of *Candida glabrata*. Cultures were incubated for 24 h at 37 ºC in the presence of copper 156 µM (laboratory) or 313 µM (clinical) of $CuSO_4$ (Cu), 32 µg/mL fluconazole (Fluc) or both (Cu+Fluc), and growth was compared to that of untreated cultures (Control). C: Checkerboard assays. *CgΔcdr1* cells were exposed to all the possible combinations of $CuSO_4$ and fluconazole (ranging between 5000 – 78 µM for $CuSO_4$ and 128 – 0.125 µg/mL for fluconazole). Growth ratios were evaluated after 24 h at 37 ºC by measuring $OD_{600}$ and normalizing it relative to untreated controls (see gradient bar). The Fractional Inhibitory Concentration (FIC) index (ΣFIC) was calculated for the indicated combinations of fluconazole and $CuSO_4$. The dashed red squares indicated the Cu+Fluc concentrations where synergy is observed. D: Tables indicating the MIC of fluconazole ($MIC_{flu}$) and the IC of $CuSO_4$ ($IC_{Cu}$) for all the strains used in this figure. MICs and ICs were evaluated after 24 h at 37 ºC with the respective compounds.

Table S1– Strains used in this study.

| Strain | Genotype/ Description | Source |
|---|---|---|
| *C.glabrata* ATCC2001 | Wild-type strain | ATCC |
| *C. glabrata* HTL | his3Δ::FRT leu2Δ::FRT trp1Δ::FRT | (1) |
| *C. glabrata* HTU | his3Δ::FRT ura3Δ::FRT trp1Δ::FRT | (2) |
| *C. glabrata* BVGC3 | ATCC2001 ERG11-3× HA | (3) |
| *C. glabrata* Δ*cdr1* | his3Δ::FRT leu2Δ::FRT trp1Δ::FRT CAGL0M01760g::NAT1 | (1) |
| *C. glabrata* Δ*zap1* | leu2Δ::FRT trp1Δ::FRT CAGL0J05060g::his3Δ | This study |
| *S. cerevisiae* BY4742 | MATα his3Δ1 leu2Δ0 lys2Δ0 uraΔ0 | Euroscarf |
| *S. cerevisiae* Δ*zap1* | MATα his3Δ1 leu2Δ0 lys2Δ0 uraΔ0 YJL056c::kanMX4 | Euroscarf |
| H17 | *Candida glabrata* clinical strain | vaginal mucosa |

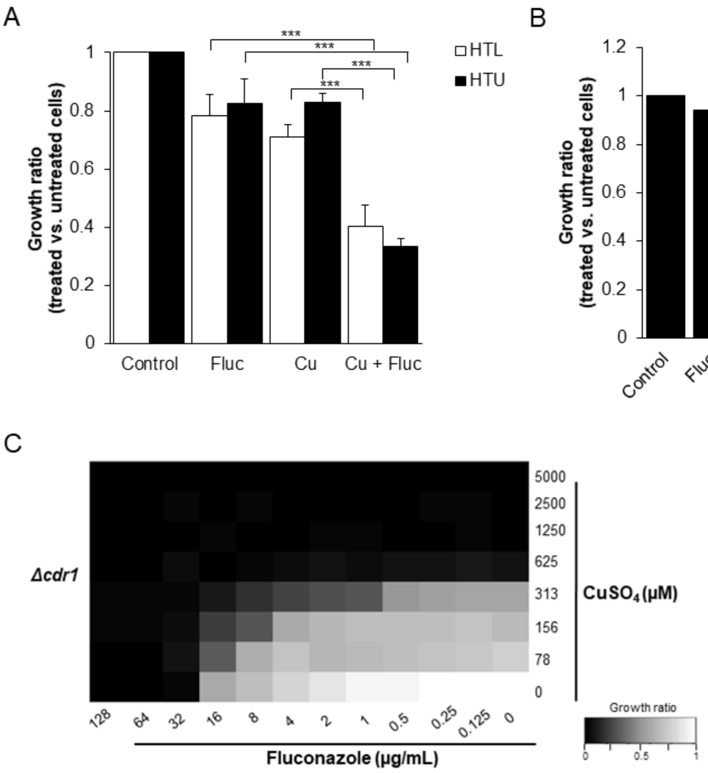


| ΣFIC | | | | | | | | | |
|---|---|---|---|---|---|---|---|---|---|
| 0.16 | 0.28 | 0.53 | 0.53 | 0.53 | 1.03 | 1.03 | 1.03 | **313** | CuSO$_4$ (μM) |
| 0.38 | 0.5 | 0.75 | 0.75 | 0.75 | 1.25 | 1.25 | 1.25 | **156** | |
| 0.63 | 0.75 | 1 | 1 | 1 | 1.5 | 1.5 | 1.5 | **78** | |
| **16** | **8** | **4** | **2** | **1** | **0.5** | **0.25** | **0.125** | | |

**Fluconazole (μg/mL)**

ΣFIC ≤ 0.5 – synergy

D

| **Strain** | **MIC$_{flu}$ (μg/mL)** Broth microdilution assay | |
|---|---|---|
| | MIC$_{50}$ | MIC$_{90}$ |
| H17 | 64-32 | >128 |
| HTU | 64-32 | 64-128 |
| HTL | 64-32 | 64-128 |
| *Δcdr1* | 32-16 | 32-16 |

| **Strain** | **IC$_{Cu}$ (μM)** Broth microdilution assay | |
|---|---|---|
| | IC$_{50}$ | IC$_{90}$ |
| H17 | 1.25-0.63 | 1.25-0.63 |
| HTU | 0.31-0.16 | 1.25 -0.63 |
| HTL | 0.63-0.31 | 1.25 -0.63 |
| *Δcdr1* | 0.63-0.31 | 0.63-0.31 |

**Figure S1** – Effect of the combination of copper and fluconazole on the growth of A: different laboratory (HTL and HTU) and B: H17 clinical strain of *Candida glabrata*. Cultures were incubated for 24 h at 37 ºC in the presence of copper 156 μM (laboratory) or 313 μM (clinical) of $CuSO_4$ (Cu), 32 μg/mL fluconazole (Fluc) or both (Cu+Fluc), and growth was compared to that of untreated cultures (Control). C: Checkerboard assays. *CgΔcdr1* cells were exposed to all the possible combinations of $CuSO_4$ and fluconazole (ranging between 5000 – 78 μM for $CuSO_4$ and 128 – 0.125 μg/mL for fluconazole). Growth ratios were evaluated after 24 h at 37 ºC by measuring $OD_{600}$ and normalizing it relative to untreated controls (see gradient bar). The Fractional Inhibitory Concentration (FIC) index (ΣFIC) was calculated for the indicated combinations of fluconazole and $CuSO_4$. The dashed red squares indicated the Cu+Fluc concentrations where synergy is observed. D: Tables indicating the MIC of fluconazole ($MIC_{flu}$) and the IC of $CuSO_4$ ($IC_{Cu}$) for all the strains used in this figure. MICs and ICs were evaluated after 24 h at 37 ºC with the respective compounds.

**References:**

1. Schwarzmüller T, Ma B, Hiller E, Istel F, Tscherner M, Brunke S, Ames L, Firon A, Green B, Cabral V, Marcet-Houben M, Jacobsen ID, Quintin J, Seider K, Frohner I, Glaser W, Jungwirth H, Bachellier-Bassi S, Chauvel M, Zeidler U, Ferrandon D, Gabaldón T, Hube B, d'Enfert C, Rupp S, Cormack B, Haynes K, Kuchler K. 2014. Systematic phenotyping of a large-scale Candida glabrata deletion collection reveals novel antifungal tolerance genes. PLoS Pathog 10:e1004211.
2. Jacobsen ID, Brunke S, Seider K, Schwarzmüller T, Firon A, d'Enfért C, Kuchler K, Hube B. 2010. Candida glabrata persistence in mice does not depend on host immunosuppression and is unaffected by fungal amino acid auxotrophy. Infect Immun 78:1066-77.
3. Vu BG, Thomas GH, Moye-Rowley WS. 2019. Evidence that Ergosterol Biosynthesis Modulates Activity of the Pdr1 Transcription Factor in Candida glabrata. mBio 10.
